# Deep integrative models for large-scale human genomics

**DOI:** 10.1101/2021.06.11.447883

**Authors:** Arnór I. Sigurdsson, David Westergaard, Ole Winther, Ole Lund, Søren Brunak, Bjarni J. Vilhjálmsson, Simon Rasmussen

## Abstract

Polygenic risk scores (PRSs) are expected to play a critical role in achieving precision medicine. Currently, PRS predictors are generally based on linear models using summary statistics, and more recently individual-level data. However, these predictors mainly capture additive relationships and are limited in data modalities they can use. Here, we developed a deep learning framework (EIR) for PRS prediction which includes a model, genome-local-net (GLN), specifically designed for large scale genomics data. The framework supports multi-task (MT) learning, automatic integration of other clinical and biochemical data, and model explainability. When applied to individual level data in the UK Biobank, we found that GLN outperformed LASSO for a wide range of diseases and in particularly autoimmune diseases. Furthermore, we show that this was likely due to modelling epistasis, and we showcase this by identifying widespread epistasis for Type 1 Diabetes. Furthermore, we trained PRS by integrating genotype, blood, urine and anthropometrics and found that this improved performance for 93% of 290 diseases and disorders considered. Finally, we found that including genotype data provided better calibrated PRS models compared to using measurements alone. EIR is available at https://github.com/arnor-sigurdsson/EIR.

## INTRODUCTION

Polygenic risk scores (PRSs) are becoming increasingly relevant to public health due to larger cohorts and the development of more powerful prediction algorithms. Today, accurate PRS predictors have been trained to predict various human diseases such as type 2 diabetes, coronary artery disease and breast cancer^1–3^. Such PRS predictors are expected to become pervasive in clinical human health and decision-making, hence playing a fundamental role in achieving personalized medicine^4–6^. PRS predictors can generally be placed in two categories based on the type of training data used, those using summary statistics from genome-wide association studies (GWAS) and those using individual-level data^7^. Today, the combined GWAS approach is more prevalent due to larger sample sizes. However, this is rapidly changing with individual-level human genetic variation data increasing in size, with cohorts comprising hundreds of thousands and even millions^8–12^. These large individual-level cohorts increasingly offer the opportunity of training accurate predictors for estimating PRSs which can outperform the combined GWAS approach^7^. Today, many established methods exist for training predictors on summary statistics^13–18^ and individual-level data^19–23^, but these predictors generally explore linear relationships.

Artificial Intelligence (AI) and Deep Learning (DL) has revolutionized several scientific fields and is changing our society. At the end of 2019 it was selected as one of ten scientific events that shaped the last decade^24^. Within life sciences, DL has gained pace within the recent years^25–27^. In particular for imaging, but also notably single-cell sequencing^28,29^, protein localization and protein folding^30,31^. For genome sequence data, the efforts have mainly been focused on identifying motifs such as ChIP-seq or identifying genomic variation^32^. Simultaneously, DL frameworks for large discrete data, such as genome-wide data, have not been extensively developed in the field. A potential advantage of DL based methods for PRS prediction is capturing complex non-linear effects, such as epistasis. However, previous work using neural networks (NNs) for predicting human traits and diseases directly from large-scale genomics has shown worse performance for NN models compared to linear ones^33,34^. The results indicate that the NNs were not able to capitalize on significant interaction effects, or that no significant interaction effects are present in the data. The latter explanation is in contrast with studies focusing on model organisms, where significant interaction effects have been found^35–37^. However, there remains both doubt and controversy regarding the role of complex interaction effects in human traits and diseases^38–42^. Therefore, there could be many explanations for why the non-linear models have not shown consistent gains over linear models in predicting human traits and diseases. These include (a) linear models are able to capture the upper bound of the genetic variance explained by genotyped SNPs, (b) for the traits tested, there are either not enough samples or not enough SNPs measured to find the interaction effects, (c) there is a significant risk of mislabelled samples in the case of diseases which could affect the model’s ability to learn subtle interaction effects, (d) interaction effects are present, but their effects are non-significant and (e) interaction effects are present and significant, but the non-linear models are not able to capture the interactions.

Despite significant non-linear interactions not having been widely shown, there are still advantages of developing NN based models for the prediction of disease risk. One factor is the flexibility of NN based models that can be developed to accept multi-modal inputs and predict more than one trait at the same time. Some of these inputs might even be unstructured and highly non-linear, such as images or text. This in itself is a clear advantage over linear models. Additionally, even though the separate modalities could mostly be modelled with linear models, there is always the possibility that complex interaction effects, e.g. between clinical measurements and other biological data exist. Using NNs allows us to discover such relationships.

However, there are many challenges with building complex NN models that can be applied to human health data. A key challenge is the immense scale of biological data. For example, genomics data often contain millions of genetic variants genotyped for large sample sizes^10,43,44^. Another challenge with fully leveraging health data is that it is often multi-modal. Supervised machine learning tasks are often trained to accept one type of input, for instance classifying the main object in a given image. By contrast, health data can be compromised of multi-omics data such as genomics, transcriptomics and proteomics data coupled with targeted biochemical and clinical data, and even include ultra-high resolution imaging. In order to provide a comprehensive disease risk assessment, methods that can account for genetic, environmental and other risk factors can be advantageous.

Therefore, we developed a DL framework that supports large-scale genomics data and can integrate it with other omics or clinical data. Among the features of the framework is a new neural network model, genome-local-net (GLN), that we specifically developed for large scale genomics data. GLN is based on a custom locally-connected layer (LCL)^45–47^ we developed, and it was able to extract genetic information from genome-wide data with equal or better performance compared to the other NN models we tested. Compared to our implementation of the least absolute shrinkage and selection operator (LASSO)^48^, we found GLN to perform statistically better overall on 338 diseases, disorders and traits. Of particular interest were autoimmune diseases such as type 1 diabetes (T1D) and rheumatoid arthritis, which previously have been shown to have complex interaction effects^49,50^. A thorough analysis of the T1D SNPs most highly activated by the GLN model shows widespread interactions between them, even across chromosomes. All models in the framework automatically extend to multi-task (MT) learning, which we showcase by training one GLN model to predict 338 diseases at the same time. Furthermore, we integrate genotype data, age and sex covariates, blood measurements, urine measurements and various anthropometrics when training GLN based models across 290 diseases in the UK Biobank (UKBB). The integration of these measurements shows a clear improvement for almost all traits, highlighting the potential of integrative models for health based predictions. Using explainable AI, we identify relevant SNPs and clinical measurements concordant with disease literature, and finally we show that integrating genotype and clinical data gives better calibrated models for precision medicine.

## RESULTS

### A locally-connected layer for genome-wide data

We initially established the ability of different candidate models to capture additive and non-linear XOR (interaction) effects when using simulated genotype data to predict a continuous output. As expected, we found that the NN-based multilayer perceptron (MLP)^51^ and convolutional neural network (CNN)^52,53^ models were able to capture and model non-linear interaction effects with *R*^2^ of 0.95-0.98. However, the linear LASSO model had *R*^2^ of 0.75 for a mix of additive and XOR effects, whereas it completely failed to model pure XOR effects with an *R*^2^ of -0.03 (Supplementary Table 1). When scaling the NN based models to genome-wide genotype data or even to whole-genome sequencing data, the number of parameters when using fully connected (FC) layers increases dramatically. For instance, an FC layer with an input of one million one-hot encoded SNPs (i.e. 4 elements per SNP) would require roughly 400 million weights to be connected to a hidden layer of 100 neurons. While convolutional layers can be much more parameter efficient, the computational complexity of training them on very high dimensional inputs can rival or exceed that of FC layers^47^. Therefore, in order to have a model that was both parameter efficient and could take advantage of the local positional variance in genomics data, we implemented a locally connected layer (Supplementary Figure 1). The layer is sparsely connected through groups, which greatly reduces the number of parameters in comparison to an FC layer. The sparse connection allows for a larger intermediate representation while still keeping the parameter count relatively low. The GLN model is composed of multiple LCLs (Figure 1a), and as was the case with the MLP and CNN models, it effectively captured both additive and non-linear effects in the simulated data (*R*^2^ = 0.98) while using fewer parameters (1.6X and 5.1X fewer than CNN and MLP respectively).

**Figure 1.**
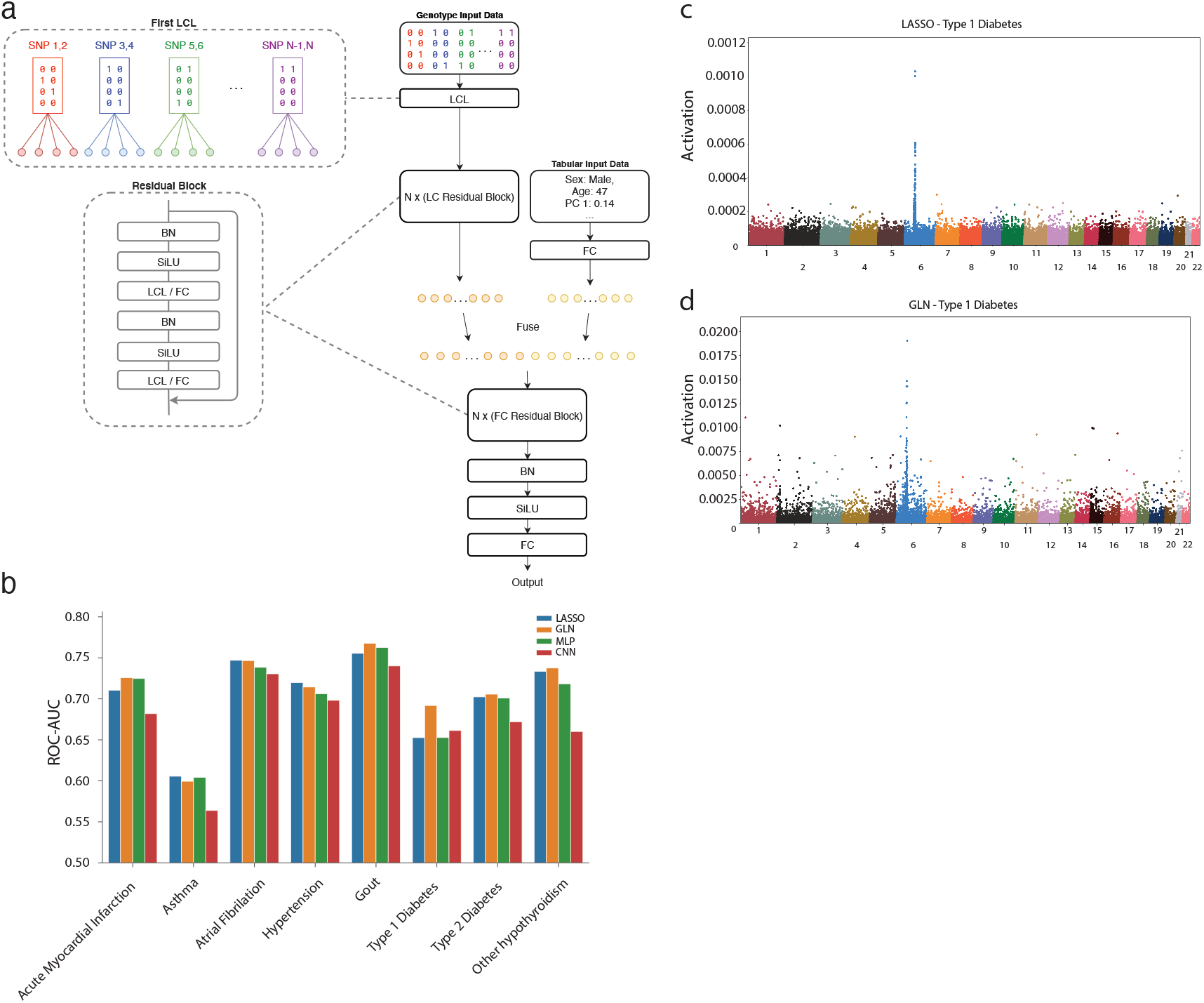
Genome-local-net (GLN) model architecture and performance. **(a)** Model architecture. The model uses a locally connected layer (LCL) with a kernel width covering two SNPs and four weight sets as the first layer. The output of the first layer subsequently goes through residual blocks composed of LCLs. The genomic representation is then fused with the tabular representation, which then is propagated through FC based residual blocks. A final set of `BN-SiLU-FC` layers is used to compute the final output. **(b)** Comparison of LASSO (blue), GLN (orange), MLP (green) and CNN (red) performance on the test set across 8 traits in AUC-ROC. All models were adjusted for age, sex and the first 10 genomic PCs. **(c)** SNP activation distribution for type 1 diabetes for LASSO, showing high activations in and around the HLA region. Each point represents a variant, and the points are colored according to chromosomes. **(d)** SNP activation distribution for type 1 diabetes for GLN, showing high activations in and around the HLA region. Each point represents a variant, and the points are colored according to chromosomes.

### GLN based Genome-wide Polygenic Scores can outperform LASSO in the UKBB cohort

We then trained and validated PRSs using either LASSO, an MLP, CNNs or the GLN model for eight selected traits on 413,736 individuals with British, Irish, or any other Western European background in the UKBB cohort (Figure 1b, Supplementary Figure 2 and Supplementary Data 1). Interestingly, we found that GLN was superior to using our LASSO model for T1D, Gout and Acute Myocardial Infarction with improvements of 0.04, 0.02 and 0.01 ROC-AUC on a held-out test set (Methods). For the remaining traits, the differences were less than 0.01 ROC-AUC. Additionally, the GLN had better performance compared to the MLP and CNN with average improvements of 0.01 and 0.04 ROC-AUC, respectively. This replicates previous results where CNN based models did not show a consistent advantage for human trait prediction^33,34^. To examine whether the gain of 0.04 ROC-AUC for T1D was due to the chosen hyperparameters for the LASSO model, we retrained the LASSO with various combinations of hyperparameters but did not find it to match the performance of the GLN model (Supplementary Table 2). To investigate and explain what the models had learned, we determined the SNPs that had the highest effects and cross-referenced them to known associations for a particular trait. Specifically, for the T1D model we found that both LASSO and GLN were activated at the HLA region of chromosome 6 (Figure 1c,d) – a region that has previously been associated with T1D^54^. However, the GLN model was also highly activated by SNPs at multiple sites across the genome (Figure 1d and Supplementary Table 3). To examine the consistency in which SNPs were highly activated by the GLN model, we aggregated the activations across 10 training runs with different seeds each (Supplementary Figure 3 and Supplementary Table 4). Interestingly, we found that among the highly activated SNPs, some were consistently found outside chr6. Examples include chr11 SNP rs3842753 located at the insulin gene (INS), which had previously been associated with T1D^55^, and chr5 SNP rs17555038, which is located in the sarcoglycan delta (SGCD) gene. Furthermore, for this variant, the main effects of the heterozygote and homozygote alternative were opposite, with the AC allele decreasing risk and CC allele increasing risk, an effect which purely additive models are not expected to capture (Supplementary Figure 4). Finally, analyzing the main effects of the top 20 SNPs identified across the training runs, we found more examples of non-additive effects for T1D (Supplementary Figure 5).

### GLN identifies disease relevant variants

When expanding the activation analysis to the other 7 traits, we found that in all cases a known association was found among the top 20 SNPs or the genes they reside in (Supplementary Figures 6–13 and Supplementary Tables 3 and 5 to 11). This is a strong indication of the models learning biologically relevant associations and that complex neural networks can be interpreted when modelled on extremely high dimensional genotype data. Even for diseases such as Acute Myocardial Infarction and Gout where the covariates (age, sex and first 10 genotype principal components (PCs)) had a better performance (Supplementary Figure 14 and Supplementary Data 2) we found that the GLN model was activated by numerous relevant SNPs and genes for both diseases (Supplementary Tables 5 and 9). The better performance of the covariate based models could be due to the covariates having much larger effects than the genotyped SNPs, e.g. if a disease was strongly affected by age or population stratification. Including the high-dimensional genotype data could increase overfitting, which then inflicts a performance trade-off against the much lower dimensionality of using only the covariates. Hence, a higher case count might be required to capture the SNP effects to such a degree that it boosts performance over the covariate based models^56^.

### Improved PRSs for autoimmune diseases

Knowing that the GLN model was competitive with the LASSO implementation on the 8 traits, we applied a more data driven approach of training GLN, LASSO and two covariate based models on 338 binary disease traits with at least 1,000 cases in the UKBB cohort (Supplementary Data 3). Among the four models tested, the GLN performed best on 59 diseases, whereas the LASSO model performed best on 42 diseases. Comparing the ROC-AUCs between GLN and LASSO, we found GLN to perform better overall (Wilcoxon signed-rank test, one-sided, *P* = 4.7 × 10^−14^). Interestingly, using only covariates had the best performance for the remaining 237 traits, and overall it performed better when compared to GLN (Wilcoxon signed-rank test, one-sided, *P* = 4.6 × 10^−15^ and *P* = 0.0015 for linear and NN based covariate models respectively) (Supplementary Data 4). The covariate based models performing better could be due to the low effective sample size (ESS) and the nature of some traits being more driven by environmental factors (Supplementary Figures 15 and 16). When filtering disease traits for where GLN and LASSO had better performance compared with covariates and difference of at least 0.01 ROC-AUC, we found 16 and 9 disease traits where GLN and LASSO had the best performance, respectively (Supplementary Figures 17 and 18). Interestingly, the GLN model performed markedly better on T1D, rheumatoid arthritis, multiple sclerosis, psoriasis and ulcerative colitis, all autoimmune traits in which studies have shown indication of interaction effects^49,50,57–61^. For instance, for rheumatoid arthritis, the GLN model had a ROC-AUC of 0.665 while the LASSO had a ROC-AUC of 0.624 on the test set and the covariate only models achieved a ROC-AUC of 0.622 and 0.634 for the LASSO and NN based models, respectively (Supplementary Figure 19). When examining GLN and LASSO activations for rheumatoid arthritis, we found, as above, the models to be activated by relevant SNPs (Supplementary Figures 20 and 21 and Supplementary Tables 13 and 14). Taken together, our results therefore show an improvement of using NNs compared to LASSO for predicting disease risk from genome-wide genomics data.

### GLN identifies SNPs with widespread interaction effects

With results showing improved performance when using GLN for traits suggested having interaction effects, we decided to analyze the T1D SNPs activated by the GLN model in more detail (see Methods). Using gradient boosted decision trees (GBDT) we identified that the strongest 200 interactions spanning 17 different chromosomes, with particularly strong effects found within chr6 but also between SNPs on chr6 and chr1, chr11 and chr12 (Figure 2a). In particular, we found the SNP rs9273363 located near HLA-DQB1 to have, as previously found, interaction with multiple other variants^62–65^. An example that we found was the interaction of rs9273363 with chr11 rs3842752 and rs3842753, which map to insulin (INS and INS-IGF2) and were in strong LD with each other. In order to examine the interaction effects more precisely, we focused on the interaction between rs3842752 and rs9273363. Individually, the AA allele of rs9273363 (HLA-DQB1) and GG allele of rs3842752 (INS) increased the model output towards a positive T1D prediction with effects of 0.6 and 0.1, respectively, whereas GA and AA of rs3842752 decreased risk (Figure 2b,c). Training a logistic regression model on the two SNPs with T1D as the target validated the direction of the main effects, with odds-ratios (ORs) of 4.36 and 1.41 for rs9273363 and rs3842752 respectively (Supplementary Table 15). The ORs were close to those from a previous T1D study for the AA allele of rs9273363 (OR 5.48) and the TT allele of another INS SNP, rs3842727 (OR 1.53), which was in high LD with rs3842752 (*R*^2^ *>* 0.75)^66^. However, when rs9273363 (HLA-DQB1) was homozygote for the risk allele (AA) the presence of at least one protective allele (GA or AA) of rs3842752 (INS and INS-IGF2) additionally decreased the risk of T1D (Figure 2d). This indicates that the GLN model was able to identify SNPs that have main and non-linear interaction effects, and that the interaction effects can be between chromosomes. Interestingly, rs9273363 tags the HLA-DQB1*03:02 allele, indicating that being homozygous for the risk allele (AA, HLA-DQB1*03:02) has a very large impact on T1D risk. Furthermore, we found that this allele had the most high T1D ranking interactions. For instance, among the top 20 SNPs interacting most strongly with rs9273363, six of them were not on chr6. Of the 14 located on chr6, 11 were not in LD with rs9273363 (*R*^2^ *<* 0.1) and besides their own main effect modified the risk contribution of rs9273363 between 0.15 to -0.3 through interaction effects (Supplementary Figure 22). Examining the output logits of the GBDT model, a value of 0.3 does have a relatively strong influence in shifting the model’s attributed risk for an individual (Supplementary Figure 23). Therefore, the total contribution of all interaction effects can have a strong influence in modulating the total risk of an individual, highlighting their importance for predictive modelling. Taken together, this provides evidence for a complex relationships between loci and alleles in modulating T1D risk that can be discovered and modeled using EIR.

**Figure 2.**
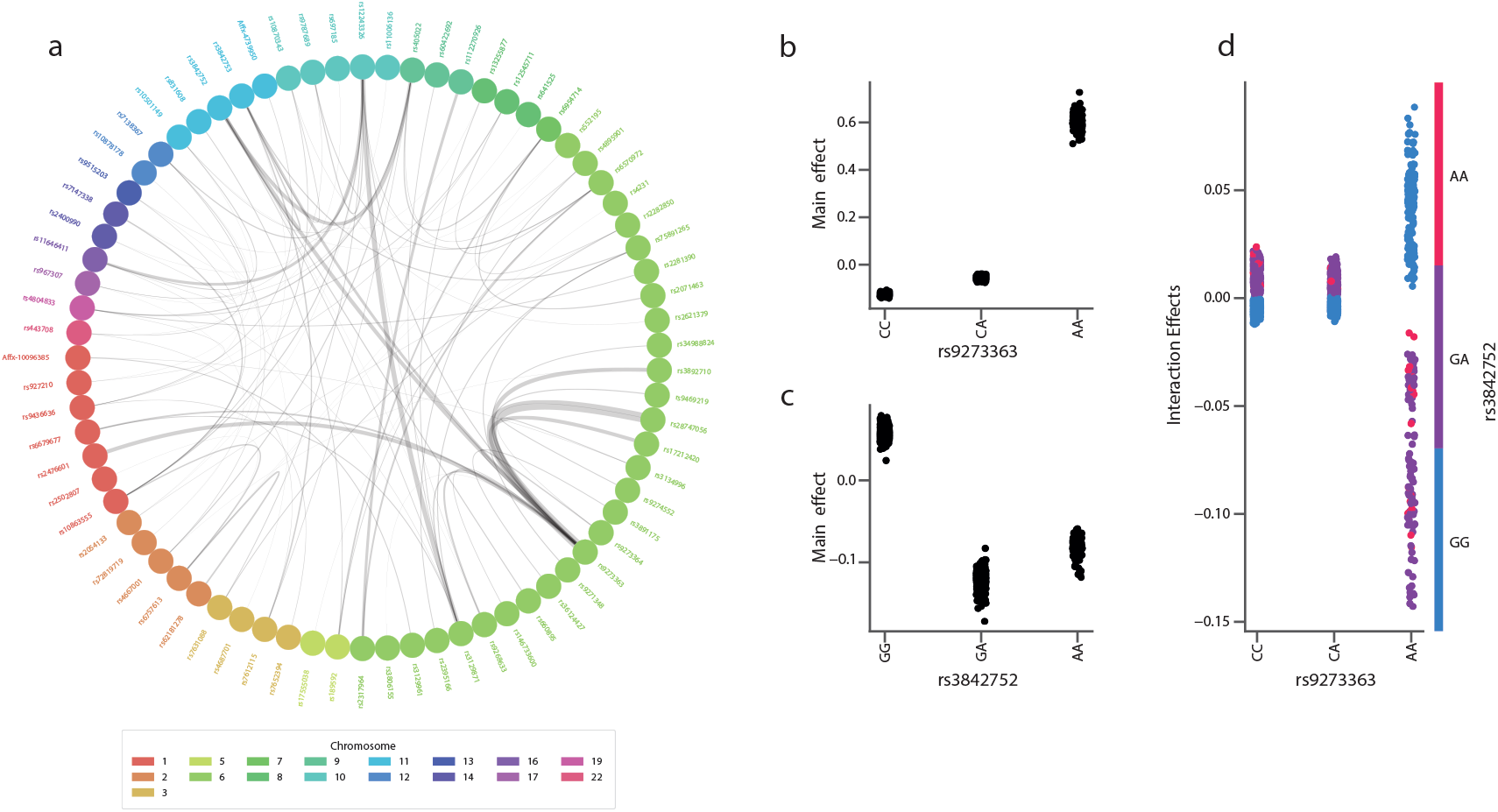
Interaction effects among highly activated SNPs for type 1 diabetes (T1D). **(a)** A network showing the interaction between different SNPs for T1D. The top 200 activated SNPs across 10 training runs with the GLN model were studied for interaction effects between them using gradient boosted decision trees (GBDT). The 200 strongest interaction effects among all SNP combinations are plotted in the graph. Each node represents a variant, and the edge widths represent the strength of the interaction between the connected variants. The node colors represent which chromosome variants reside in. The network shows particularly strong interaction effects between various SNPs on chr6, but also widespread interaction effects between chromosomes. **(b)** Main effects of chr6 SNP rs9273363 on T1D, with the AA allele having a strong effect in increasing risk. The y-axis values represent the main effect influence of a given rs9273363 allele on the trained GBDT model output logits. **(c)** Main effects of chr11 SNP rs3842752 on T1D, with the GG allele having a moderate effect in increasing T1D risk. The y-axis values represent the main effect influence of a given rs3842752 allele on the trained GBDT model output logits. **(d)** Interaction effects between chr6 SNP rs9273363 and chr11 SNP rs3842752. The x-axis represents the rs9273363 allele, the y-axis represents the interaction effect influence on the trained GBDT model output logits and the colors represent the GG (blue), GA (purple) and AA (red) alleles of rs3842752. The vertical dispersion seen for the AA allele of rs9273363 indicates that allele combinations explored have different effects for different samples. This can be due to other SNPs having an additional interaction effect on rs9273363 and rs3842752, which can be seen in figure 2a where the SNPs not only interact with each other, but multiple other SNPs.

### GLN is fast and robust to missing data

To simplify calculation of PRSs we implemented the models, including the LASSO, to automatically handle missing genotype data and thus removing the need to impute data prior to training. In this work, the genotype data was not pre-processed extensively prior to modelling (denoted “NO-QC”). To investigate whether our results were consistent when using traditional pre-processing (denoted “QC”), we also trained GLN and LASSO on QC data. Besides reducing the number of SNPs and samples considered, the QC approach additionally results in a different train/test split. The NO-QC approach gave slightly better results on our eight benchmark traits (Supplementary Figure 24 and Supplementary Data 5). However, we did see the overall trends remain consistent whether using QC or NO-QC, e.g. with GLN performing markedly better on T1D (Supplementary Figure 24). Finally, we investigated the computational costs of training the PRS models. For the eight benchmark traits, we found training the GLN to be slightly faster (32 hours) compared with our LASSO (34 hours) (Supplementary Figure 25 and Supplementary Data 6). Even though the training latency of the LASSO model was lower than any of the NN based models, the total training time was higher due to using more steps before model convergence. Therefore, the framework was able to train large and deep neural networks on high dimensional individual-level genotype data in a reasonable time.

### Multi-task learning offers a trade-off between performance and complexity

In multi-task (MT) learning, a single model is trained to solve multiple objectives at the same time, such as predicting height, disease liability and ethnicity. This can lead to improved predictive performance, reduced training time and better parameter efficiency^67,68^. We therefore hypothesized that predicting multiple outcomes simultaneously could regularize and potentially improve prediction performance. Using Type 2 Diabetes (T2D) for comparison, we trained MT models to predict two, eight and 338 diseases jointly and found that maximum validation performance got progressively worse when increasing the number of tasks (Figure 3a and Supplementary Data 7). This indicates that the model capacity was not high enough to effectively capture the variance of multiple traits as well as the single task model, or that negative transfer between tasks degraded performance^69^. Similarly, when comparing test set performance for the respective single task models and an MT model trained on the 8 benchmark traits, we found that the MT model was slightly worse for 7 diseases (average 0.023 ROC-AUC lower), with Acute Myocardial Infarction being the exception (0.0054 ROC-AUC improvement) (Figure 3b). However, despite being slightly worse for most of the traits, the MT models were remarkably effective. For example, the 8 trait MT model had a test ROC-AUC of 0.68 for T1D, which was considerably higher than the 0.58-0.59 ROC-AUC when using only covariates. In order to examine how well the framework scaled and whether we could effectively train very large scale MT models, we trained one GLN model to jointly predict 338 traits simultaneously. As expected, modelling on all traits jointly significantly reduced the training time (11X) and number of parameters per trait (395X) (Figure 3c,d). As in the other MT experiments, this came at the cost of reduced performance compared to the single task setting (Wilcoxon signed-rank test, one-sided, *P* = 0.025), however only with an average difference of 0.0056 ROC-AUC (Figure 3e and Supplementary Data 8). Compared to the best performing covariate based models for each trait, the large MT model performed better for 61 traits, indicating that it was able to effectively capture genotype variance for some traits and not only using the covariates.

**Figure 3.**
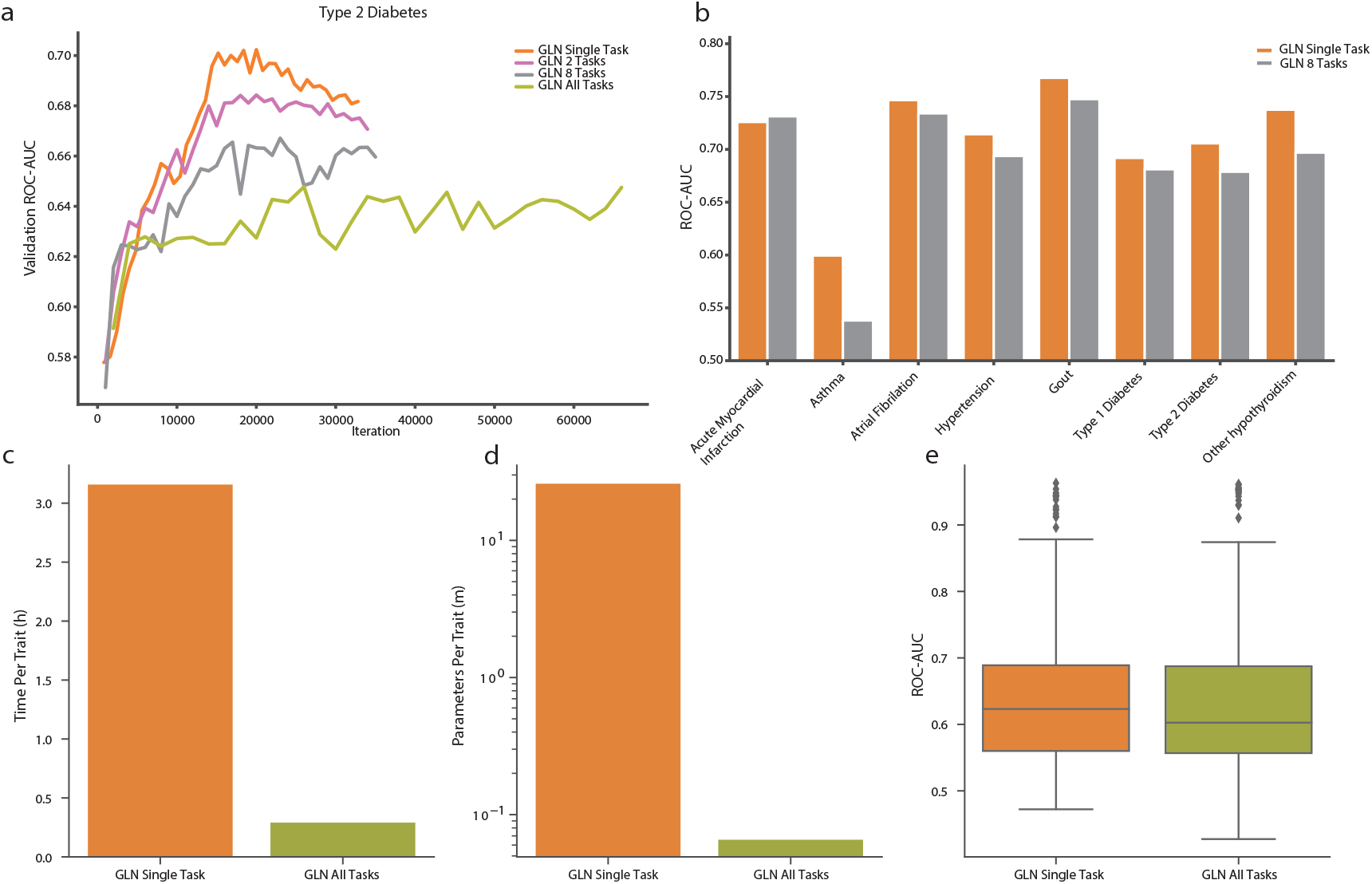
Genome-local-net (GLN) multi-task (MT) predictions. **(a)** Comparison of validation curves in ROC-AUC for Type 2 Diabetes (T2D) as more tasks are added alongside the single task type 2 diabetes prediction (orange). The two task model (pink) is trained jointly on T2D and hypertension. The 8 task model (gray) is trained on the 8 benchmark traits showed in figure 1b. The model trained on all tasks (green) is trained on the total set of 338 diseases considered in this work. The single and two task runs show signs of overfitting, as validation performance peaks and starts to deteriorate around 20K iterations. The eight and all MT runs do not show as clear signs of overfitting, but overall performance is worse. **(b)** Comparison of single task (orange) and MT performance (gray) for the 8 benchmark traits on the held-out test set. **(c)** Comparison of training time per trait for the all task MT model (green) and single task training (orange). **(d)** Comparison of number of parameters per trait for the all task MT model (green) and single task (orange) training. **(e)** Overall performance on the held-out test set of the all task GLN MT model (green) and single task (orange) training.

### Integrating genomics with clinical data improves predictive performance

Although genetic data has proved to be a powerful predictor of various traits and diseases, there are other factors such as environmental effects that can play an important part^70^. With the increased digitization in the healthcare industry, clinical and electronic health data is only expected to become more widely available. Among these are factors that are relatively easy and non-invasive to measure, such as anthropometrics, and other measurements included in the UKBB such as blood and urine measurements. To examine the benefit of using these measurements as part of our models for computing PRSs, we trained GLN models using only genotype data and covariates, and compared this with using genotype, covariates, physical, blood and urine sample measurements (denoted “Integrated”). Furthermore, to minimize data leakage we removed samples where the diagnosis occurred in the past compared to the measurements, ensuring that the model predicted future diagnoses and not previously diagnosed conditions. Therefore, when including the measurements the number of cases was for most traits reduced, leading to a trade-off between the gain of including measurements and the loss of removing samples. To examine this trade-off more precisely, we compared to two genotype datasets, one where the censored individuals were removed (denoted “Genotype Filtered”), and another set where all individuals were included (denoted “Genotype”) (Figure 4a and Supplementary Data 9). For all eight benchmark traits, as expected, removing samples reduced performance with ROC-AUC of 0.014-0.092 (Wilcoxon signed-rank test, one-sided, *P* = 0.0039). Another contributing factor is that the sample removal is likely biased towards those individuals that have a high genetic load, and therefore diagnosed early. Compared to Genotype Filtered data, we found that using Integrated data greatly improved performance, with ROC-AUC increasing by 0.043-0.27 (Wilcoxon signed-rank test, one-sided, *P* = 0.0039) (Figure 4a). This was also the case when using Matthews correlation coefficient (MCC) as metric, which improved between 0.010-0.35 (Supplementary Figure 26). However, compared to the unfiltered Genotype data, the results were more disease dependent. For instance, filtering hypothyroidism for time of diagnosis reduced case count from 16,894 to 4,663 in the training set, which was reflected in ROC-AUC performance reduction of 0.092. Including measurements therefore did not outweigh the performance reduction of discarding cases. Interestingly, we found that using Integrated data had superior ROC-AUCs for five of the traits compared to using measurements and covariates only (denoted “Measurements”), highlighting the benefit of including genotype data.

**Figure 4.**
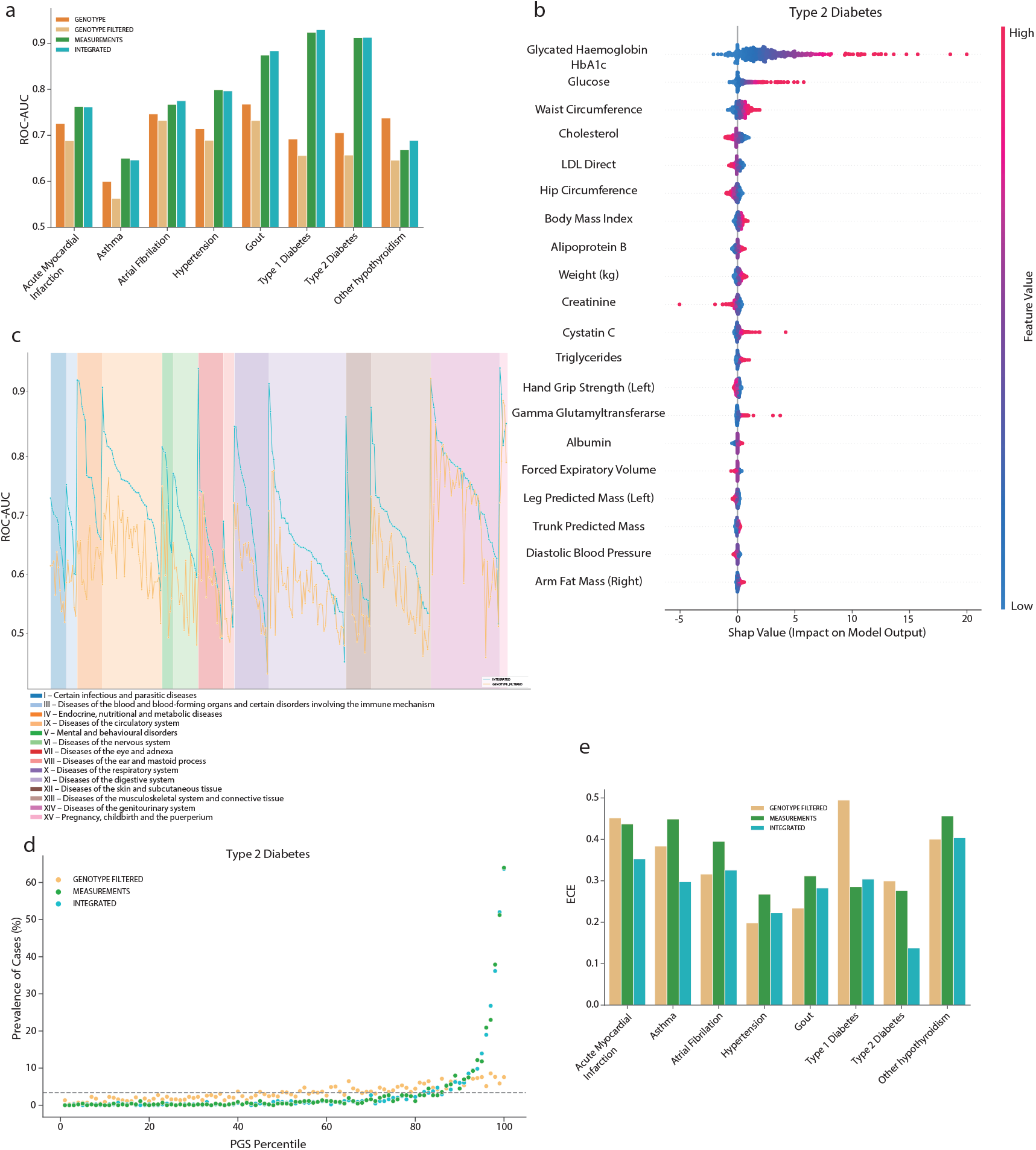
Integrating genotype and clinical data with genome-local-net (GLN). **(a)** Comparison of model performance using Genotype (orange), Genotype Filtered (light orange), Measurement (green) and Integrated (teal) data in ROC-AUC on the held-out test set. **(b)** Feature importance and impact of integration measurement values on the GLN model prediction for T1D. For a given feature, each dot represents a sample in the test set. The colors indicate the actual feature value. For example, a strong trend of high glycated haemoglobin value influencing the model to make a positive prediction for T2D can be seen. **(c)** Summary of ROC-AUC performance on the held-out test set across all the 290 traits that had a time measured column associated with them, with Integrated data (teal) compared with Genotype Filtered data (light orange), filtered for time of diagnosis. The different background colors represent different ICD-10 chapters. **(d)** Prevalence plot of T2D prediction on the held-out test set when using the GLN model on Genotype filtered (light orange), Measurement (green) and Integrated (teal) data. **(e)** Comparison of Expected Calibration Error (ECE) on the held-out test set when using the GLN model on Genotype filtered (light orange), Measurement (green) and Integrated (teal) data

### Integration of clinical and genomics data improve prediction of T2D

When investigating the activations of the integrative predictor, we found that the model was activated by relevant clinical measurement features such as glycated haemoglobin (HbA1c) and blood glucose for T2D (Figure 4b). However, for some diseases, such as T2D, predictors using Measurements data and the Integrated data had very similar ROC-AUC performances. This does not necessarily indicate that the genetic component of the traits was low, perhaps the more likely explanation is that the measurements can act as a proxy for the genomics effects. For example, in the case of T2D, high genomics risk will in many cases manifest itself in high levels of glycated haemoglobin, and when it can be measured directly, there is perhaps not much extra variance gained when including the genotype data. However, when we investigated the genotype activations, we found that the model was activated by relevant SNPs even when measurement data was included (Supplementary Figure 27). This was a similar trend we observed above, where the addition of genotype data resulted in worse overall performance, despite the models being activated by disease relevant SNPs. When exploring this further, we found that including the genotype data resulted in a predictor for T2D with higher MCC (0.43) compared with using only the measurements (0.33) (Supplementary Figure 26). Interestingly, this was particularly due to better classification of true negatives and indicates the usefulness of the integration.

### Large scale integrative modeling

Knowing that the integrative models could increase performance and learn relevant features, we, as before, performed large scale analysis of 290 traits that included time of diagnosis. Integration of the measurements showed a large increase in performance for almost all the traits compared with the Genotype Filtered predictor (Figure 4c and Supplementary Data 10). Interestingly, we had expected improvements for endocrine, nutritional and metabolic diseases but found improvements in ROC-AUC and MCC for other classes of diseases such as mental and behavioral disorders (Supplementary Figure 28). Compared with the Genotype predictors we observed the same overall trend that including measurements improved performance, but the effect was less pronounced due to the Genotype models using more samples (Wilcoxon signed-rank test, one-sided, *P* = 6.1 × 10^−37^ and *P* = 9.9 × 10^−09^ for ROC-AUC and MCC, respectively) (Supplementary Figures 29, 30 and Supplementary Data 11). To examine the effect of including genotype data when measurements were available, we compared the Measurements based models to models using Integrated data and found that the differences was small for most traits (Wilcoxon signed-rank test, one-sided, *P* = 0.048 and *P* = 0.054 for ROC-AUC and MCC, respectively) (Supplementary Figures 31, 32 and Supplementary Data 12). This could be due to low ESS for many of the traits, traits being driven more by environmental effects and/or high genomic risk being reflected in the measurements (Supplementary Figures 15, 16 and 33).

### Genotype data calibrates PRS from measurement data

For clinical decision-making, performance measurements such as the ROC-AUC and MCC are often insufficient to describe model performance^71,72^. We therefore examined the risk score distribution of our models, specifically investigating the relationship between the fraction of positives for a given percentile of the PGSs (Figure 4d and Supplementary Figure 34). When using Integrated data, we found that 5 of the 8 diseases had a prevalence of 10% or more in the top percentile, whereas models using Genotype Filtered or Measurements data had the same performance for 2 and 5 diseases, respectively. However, we also found that the models were poorly calibrated due to overconfidence, an issue that could be amplified by mislabeled data, where individuals have a high risk but have not yet been diagnosed with a disease (Supplementary Figure 35). However, for 7 of the 8 traits, including T2D, we found the Integrated model to perform better than the Measurement model in terms of Expected Calibration Error (ECE) and Brier Score (BS) (Wilcoxon signed-rank test, one-sided, *P* = 0.0078 for both metrics) (Figure 4e), indicating more reliable probability estimates, the probability estimates being closer to the true labels and better uncertainty estimation. These results are particularly interesting for traits like Acute Myocardial Infarction, where the Integrated models and Measurement models had virtually the same ROC-AUC and MCC, but a 0.084 and 0.066 improvement in ECE and BS, respectively, when including genotype data (Figure 4e and Supplementary Figure 36). These results showcase the clinical potential of integrating genotype data with clinical measurements.

## DISCUSSION

Here, by developing DL models specifically for large scale individual-level genotype data, we show that they can achieve competitive performance for a wide range of diseases. Particularly, the GLN model outperformed our LASSO implementation for various autoimmune diseases and was able to identify interaction effects. By focusing on the explainability of our GLN model, we confirmed that it could identify disease relevant variants, showcasing that complex DL models are not strictly uninterpretable black boxes. Furthermore, our model could identify widespread interaction effects. While interaction effects are often overlooked in PGS studies, we found them cumulatively to have surprisingly strong effects in some cases, e.g. 52% of the total effect of rs9273363 on T1D risk. Additionally, they can provide valuable biological insights and accounting for them provides better predictive performance for some traits. We expect that finding such complex effects will become more common in the future, especially with the development of larger, better phenotyped cohorts. Interpreting such associations should be done with care, however, as computational associations are not guaranteed to capture true biological effects^73^. Furthermore, while complex non-linear models can be used to uncover such effects and provide a relative comparison of their strength, once identified, linear methods could be used to explicitly model and quantify the effects. Additionally, we did not filter for LD which can dilute the activation signal across multiple SNPs, instead of being concentrated to one SNP representative of the LD block. Furthermore, we showcase the flexibility that DL architectures offer by training a single model to predict 338 disease traits at the same time with minimal loss in performance. An interesting avenue of research could be to examine MT learning with focus on related tasks and applying more recently developed MT learning NN architectures, which might yield better results compared to our approach.

Additionally, given that a predictor that uses additional measurements, such as blood, urine and anthropometrics, can be trained on the same individuals we found a clear advantage for the case of integration. However, if including the measurements poses a data leakage risk and subsequent removal of samples, one must consider whether the trade-off in samples and additional features is acceptable. Nonetheless, we saw a strong indication that inclusion of measurements outweighed the removal of samples for many disorders. Finally, we found that the inclusion of genotype data in addition to clinical and biochemical measurements provided better calibrated models, which is highly relevant for clinical decision-making. The overall poor calibration, however, highlights the importance of examining it, especially when developing models for clinical applications. This is particularly important when using NNs, which tend to be overconfident^74^. A potential solution can be to calibrate predictions post-hoc with methods such as Platt scaling, temperature scaling or isotonic regression^75–77^. However, additional factors to consider will be uncertainty estimation and out-of-distribution detection, which post-hoc calibration methods are not guaranteed to solve.

We only considered data from individual-level cohorts, but it is straightforward to integrate PRSs from predictors trained using summary statistics or genome-wide data. Addition of these could potentially improve performance even further. Furthermore, an additional limitation of this study is that we only considered Western European ancestry. Developing predictive models that perform robustly across multiple ancestries will be of great importance. Although some attempts have been made at solving this problem with linear models^78^, examining NN based model performance in this context might provide valuable insights and results. Finally, we only considered two input modalities, genotype and tabular data. However, this only accounts for a fraction of the types of health data commonly measured, with other examples including high resolution imaging, multi-omics and electronic health data. We believe that the development of accurate predictors that can model on various types of data, whether structured or unstructured, will be important for achieving precision medicine in the future.

## METHODS AND MATERIALS

### Processing of UK Biobank genotype and clinical data

The genotype data was processed using Plink^79^, version v1.90b6.10. After processing, the genotype data was converted to 459,576 one-hot encoded sample arrays of shape (4, 803,113) each, in the case of no quality control (NO-QC). The NO-QC approach might include signals from rare variants that otherwise would be filtered out using a minor allele frequency threshold, and previous work have shown negligible differences between kinship filtered and unfiltered sets of the UKBB data^80^. When applying quality control (QC), we used the following parameters in Plink `--maf 0.001 --geno 0.03 –mind 0.1` as well as removing samples with a kinship of more than 0.1. After applying QC, there were 425,439 one-hot encoded sample arrays of shape (4, 662,143) each. X chromosomes were included in both cases. Unless otherwise specified, age, sex and the first 10 genotype principal components were included during training. For the tabular data, continuous columns were standardized using the training set statistics in all experiments, meaning that the values computed for the training set were applied to the validation and test sets. Missing values were imputed with the averages from the training set. Categorical columns were numerically encoded, missing values were marked as “NA” prior to numerical encoding. ICD-10 codes were used to derive the disease phenotypes. Only samples with a self-reported British, Irish, or any other Western European background were used for the experiments, which amounted to 413,736 samples in the training/validation sets and 45,840 in the test set in the NO-QC case. In the QC case, this resulted in 382,894 and 42,545 samples in the training/validation and test sets respectively. For the integration experiments, we removed samples where the measurements used for integration were measured after a disease diagnosis. This was to avoid feature leakage, where e.g. a drug for a certain disease influences measurement values. An alternative could be to mark the measurements as missing and allow them to be subsequently imputed with the train set statistics. However, this might bias the case data towards having all the measurements imputed, which the model might learn. Hence, it is not certain that such approaches would completely prevent feature leakage. Out of the 338 diseases, 48 did not have any time of diagnosis associated with them, and we therefore excluded these from the integration experiments and analysis.

### Training implementation and approach

All models were implemented using Pytorch^81^, version 1.7.1. A held-out test set was used for all models in order to get a final performance after training and evaluating on train and validation sets, respectively. We used negative log likelihood loss during training for the classification tasks. All models were trained with a batch size of 64 except for the large MT model (338 traits) which used a batch size of 32. During training, we used plateau learning rate scheduling to reduce the learning rate by a factor of 0.2 if the validation performance had not improved for 10 steps. The validation interval was calculated dynamically based on the number of cases for a given disease trait (*C/B* where *C* is case count and *B* is batch size, with thresholds of 100 and 2000 for the minimum and maximum intervals respectively), as was the number of validation samples used (max[10000, -1.5 = *C* + 50000] where *C* is case count). We used early stopping to terminate training when performance had not improved for a certain number of validation steps. We used 16 and 20 steps for traits with less than and more than or equal to 2,500 cases, respectively. For the early stopping, we also used a buffer of a certain number of iterations before it was activated, using 1,000 iterations for the 8 trait benchmark and the MT experiments and 2,000 iterations for the large scale experiments. Weighted sampling with respect to the target variable was used in all runs during training. All models were trained with the Adam optimizer^82^. In the NN based models, we used a weight decay of 1 × 10^−3^ with decoupled weight decay regularization^83^. All NN based models used a learning rate of 1 × 10^−4^, while the LASSO models used a learning rate of 5 ×10^−5^. We found that lower learning rate for LASSO gave better training stability and overall results. All neural network architectures used the SiLU^84,85^ (also known as Swish^86^) activation function with a trainable parameter *b* inside the sigmoid function. When using weight decay, we did not apply it to the *b* parameter. For the neural network models, we augmented the input by randomly setting 40% of the SNPs as missing in the one-hot encoded array, this is similar to input dropout^87^ and we found it to be important to prevent overfitting in the NN models. For the LASSO, we used L1 regularization with *λ* = 1 ×10^−3^ for traits that had more than 2,500 cases and *λ* = 1 ×10^−2^ for traits that had less. All models were trained on a single 16GB NVIDIA® V100 Tensor Core GPU.

### Architectures

This section details how the model architectures were implemented, which are broadly depicted in Supplementary Figures 37 and 38. The MLP feature extractor was one FC layer with 10 output nodes. The main building blocks of the convolutional feature extractor were residual blocks, with the first block using full pre-activation^88,89^. We added squeeze-and-excitation (SE) blocks^90^ to the residual blocks, which we found both stabilized training and improved performance with minimal computational overhead. We used a dropout^87^ of 0.5 between the convolutional layers in the residual blocks, as recommended in prior work^91^. Before the residual blocks, the feature extractor used a single convolutional layer with a kernel size of (4, 39), a stride of (1, 10) along and 64 output channels. All the residual blocks used 64 input and output channels, a kernel size of either (1, 20) or (1, 19) and a stride of (1, 10) in the first convolutional layer and when downsampling the identity. The feature size after the convolutional blocks was 576, which went through `BN-ACT-FC` layers with an output feature size of 256. The feature extractor of the GLN model was similar to that of the CNN model, where the main difference was that we used LCLs instead of convolutional layers, only two residual blocks instead of four and no SE blocks. In the first LCL, we used a kernel width of 8 (covering two SNPs per group) and 4 output sets and in the subsequent residual layers, we used a larger kernel width of 32 and 4 output sets. The final output dimension from the feature extractor was 396. The tabular feature extractor used in all models used embeddings for categorical inputs and left continuous inputs unchanged. The tabular inputs were concatenated and passed through a single FC layer. The fusion model aggregated the intermediate representations from the individual feature extractors by simply concatenating them. For the CNN and GLN NN predictors, we used the fused features from the fusion module as input and propagated them through FC residual blocks. For the CNN and GLN models, the predictors used four residual blocks with 256 nodes in the FC layers and a dropout of 0.5 between the FC layers. After the final residual blocks there was a `BN-ACT-DO-FC` which computed the final output for a given task. In the MLP case, we did not use residual blocks, but rather a classic feed forward network. The intermediate representation from the fusion model was propagated thorough 5 sets of `BN-ACT-DO-FC` layers. Excluding the last, all FC layers had 256 output nodes. We used a dropout of 0.5 before the FC hidden layers.

### Multi-task prediction

For each task, the NN predictor was a sequence of 4 residual blocks with FC layers composed of 256 nodes in the two and eight task models, but 64 nodes in the 338 task model. We used homoscedastic uncertainty to weight the loss contributions from individual tasks^92^. The technique we use for our MT learning is known in the as hard-parameter sharing, where all tasks share a subset of the model parameters throughout the entire training procedure. To examine how well the default GLN model performed in MT learning compared to other NN models, we compared it with an MLP model and a GLN based model using a Multi-gate Mixture-of-Experts (MGMoE)^93^ as the predictor on the 8 benchmark traits. We found the default GLN model to perform the best overall (Supplementary Figure 39 and Supplementary Data 13).

### Main and interaction effect identification

To examine the effects between SNPs, we used the 200 most highly activated SNPs by the GLN model as candidates for the analysis. Using those SNPs as inputs, we trained a gradient boosted decision trees model using the XGBoost framework^94^. The trained decision trees used a learning rate of 0.002, maximum depth of 6, 10,000 boosting iterations and a 50% training set subsample for each boosting iteration. The same training, validation and tests sets were used as for the GLN model training and evaluation. After training, we subsampled a maximum of 2,000 samples per class in the test set for the main and interaction effect analysis, for which we computed the SHAP effect values for analysis.

### Framework

We present a framework that offers training with both linear models and various DL architectures on one-hot encoded omics and tabular input data, with the option of training on both continuous and categorical target variables. These models include linear and logistic regression models, MLP, CNN and GLN. Note that as the data is encoded in a one-hot format, the linear models should be able to detect some non-additive effects such as dominance. When modelling with different datasets of omics data, it is very common for datasets to differ in the number of genotyped SNPs. This can present a challenge when applying NN models designed for one dataset to another, as the network might have to be reconfigured to match the genotype dimensionality of the new dataset. To reduce the manual work that comes with such reconfiguration, the architecture of all models in the framework is automatically configured according to the size of the omics data so that no changes are needed whether the model has, for instance, 2,000 or 803,113 genetic variants as input. The framework attempts to strike a good balance between usability and flexibility by offering various model configurations that work with the automated architecture setup. For example, in the case of CNN models, the number of channels, number of layers, kernel width, down stride and other hyperparameters can be configured when setting up the model. Users are not constricted to the predefined models however, and can when using the Python interface define their own models which can be passed to the framework. Besides configuring model architecture, various other settings are available. For learning rate scheduling, the model offers various schedules such as cosine annealing with and without warm restarts and plateau scheduling. Additionally, the number of warm-up steps can be configured. When training predictive models on extremely high dimensional data, care must be taken to avoid overfitting. This is especially true when working with complex DL models, and hence the framework offers various regularization strategies. These include weight decay, L1 regularization, dropout and input dropout for omics data. Additionally, the framework offers data augmentation strategies in the form of Mixup^95^ and two CutMix^96^ variants, which are applied both to omics and tabular data when used. We found the mixing augmentations strategies as a promising way to reduce overfitting and increasing model generalization performance when applied to omics and tabular data (Supplementary Figure 40 and Supplementary Data 14). In addition, the framework offers weighted sampling with respect to arbitrary columns in the label file, which we found to greatly speed up training when applied to the target labels. The framework automatically extends to multi-task settings when more than one target is used for the modelling task. Additionally, the framework automatically combines information from omics and tabular data when using both as inputs. To gain insights into what the models learn during training, the framework offers automated activation computation and analysis for both omics and tabular inputs, using the SHAP framework^97^. We confirmed the models learning disease relevant variants and genes using LitVar and DisGeNet^98,99^. Lastly, the framework offers early stopping and saving of the model state that performed best on the validation set across the whole training run. The functionality to use an exported model to predict on a test set or other new samples is included in the framework.

## ACKNOWLEDGMENTS

We thank Ingi Thor Sigurdsson for helpful discussion and contribution to the software. This research has been conducted using the UK Biobank Resource application 31823. A.I.S., D.W., S.R. and S.B. were supported by the Novo Nordisk Foundation (grants NNF18SA0034956, NNF14CC0001, and NNF17OC0027594). B.J.V. was supported by a Lundbeck Foundation Fellowship (R335-2019-2339) and by the Danish National Research Foundation (Niels Bohr Professorship to Prof. John McGrath), and by the Lundbeck Foundation Initiative for Integrative Psychiatric Research, iPSYCH (R102-A9118, R155-2014-1724 and R248-2017-2003). O.W was supported by the Novo Nordisk Foundation through the Center for Basic Machine Learning Research in Life Science (grant no. NNF20OC0062606). Computations described in this paper were performed using the National Life Science Supercomputing Center - Computerome at DTU and UCPH, www.computerome.dk.

## AUTHOR CONTRIBUTIONS

S.R. conceived the study and guided the analysis with D.W., S.B., O.W., O.L. and B.J.V. A.I.S. wrote the software and performed the analyses. A.I.S, B.J.V. and S.R. wrote the manuscript with contributions from all coauthors. All authors read and approved the final version of the manuscript.

## CODE AVAILABILITY

The code and binaries for EIR are available under the AGPL-v3 license and can be found at https://github.com/arnor-sigurdsson/EIR and https://github.com/RasmussenLab/EIR.

