## Supplementary Information for "Deep integrative models for large-scale human genomics"

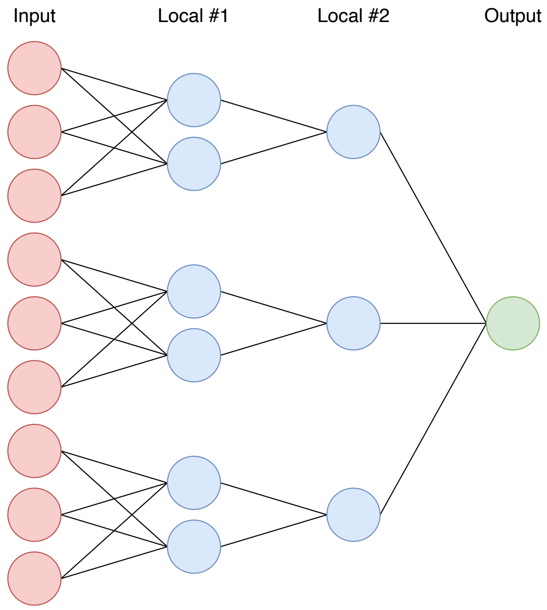


**Supplementary Figure 1.** Diagram showing how the locally-connected-layers (LCLs) are structured. The input (red) is connected to the first LCL layer, which has a kernel width of three in the first layer and two output sets, resulting in an intermediary representation with 6 nodes (light blue). The second LCL layer has a kernel width of two with one output set, resulting in an intermediary representation of 3 nodes (light blue). The intermediate representations from the second LCL go through an FC layer to generate the final output (green).

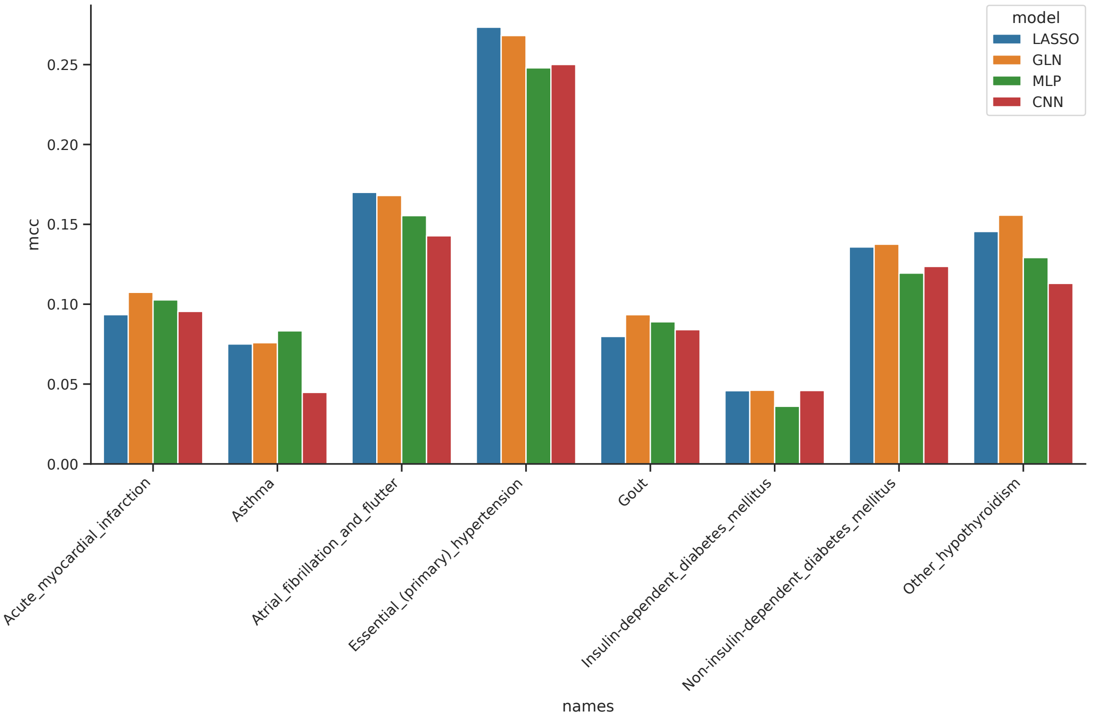


**Supplementary Figure 2.** Comparison of LASSO (blue), GLN (orange), MLP (green) and CNN (red) performance on the held-out test set across 8 traits reported in Matthews correlation coefficient (MCC). All models were adjusted for age, sex and the first 10 genomic principal components (PCs). It should be noted that the model checkpoint used was the best performing one on the validation set using the ROC-AUC. Hence, there might not be a full correlation between the chosen model checkpoint being the one that performs best with respect to MCC.

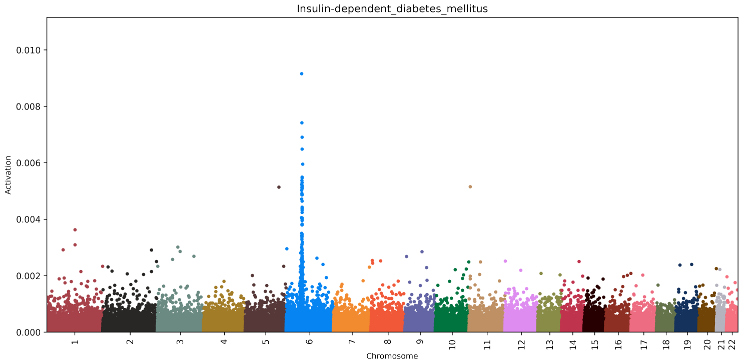


**(a)** Aggregated.

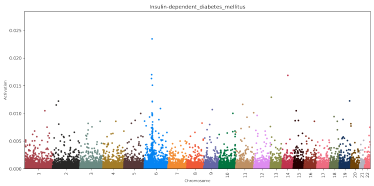


**(b)** Run 1.

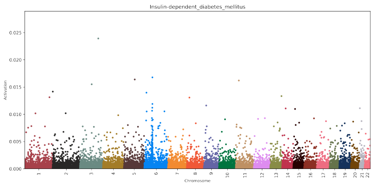


**(c)** Run 2.

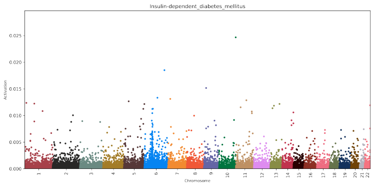


**(d)** Run 3.

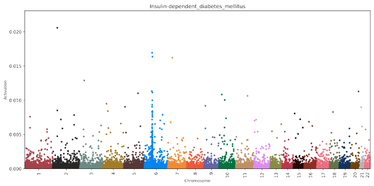


**(e)** Run 4.

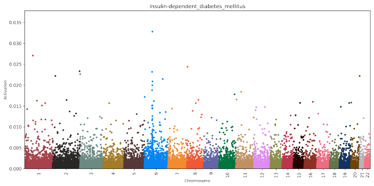


**(f)** Run 5.

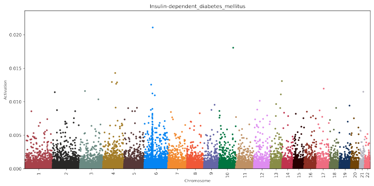


**(g)** Run 6.

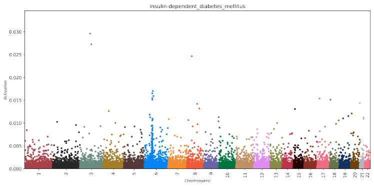


**(h)** Run 7.

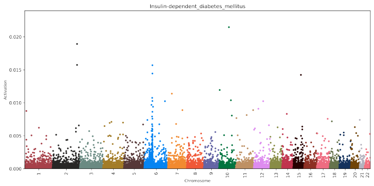


**(i)** Run 8.

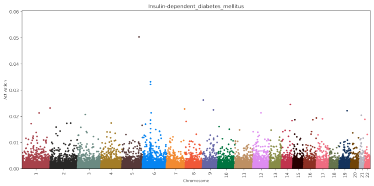


**(j)** Run 9.

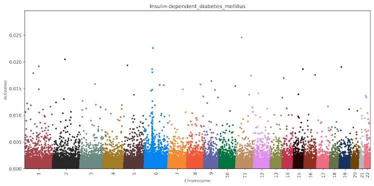


**(k)** Run 10.

**Supplementary Figure 3.** Aggregated and individual activations for type 1 diabetes (T1D) across 10 training runs with different random seeds when using the genome-local-net (GLN) model. The values on the y-axis represent a given SNP's absolute influence on the model's raw output score (logit) for T1D. While major patterns can be identified in the single runs, such as high activation in the HLA region, there is some variance in which SNPs are highly activated in other parts of the genome, where linkage disequilibrium (LD) and stochasticity during model training can contribute to the variance. By averaging the activations across single runs (aggregated), a clearer picture of which SNPs play the most important part can emerge.

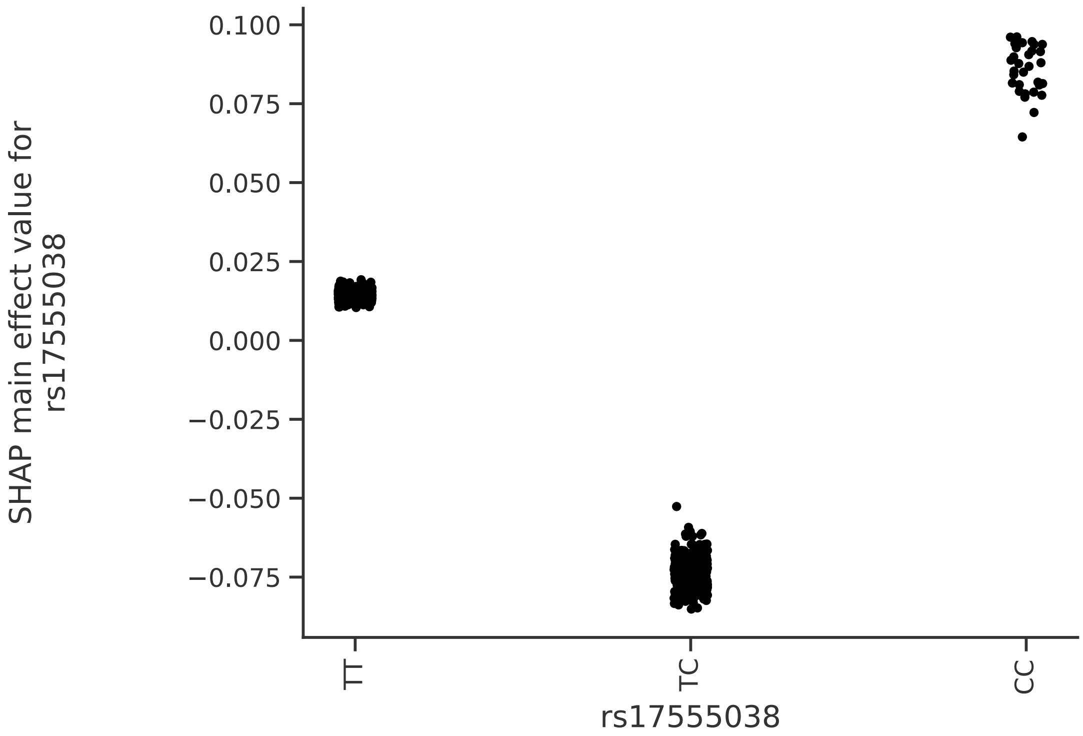


**Supplementary Figure 4.** Analysis of chr5 rs79213347 main effect on Type 1 diabetes (T1D) risk. The heterozygote TC allele decreases risk, while the homozygote CC allele increases risk. The y-axis values represent the main effect influence of a given rs79213347 allele on the trained gradient boosted decision tree (GBDT) model output logits.

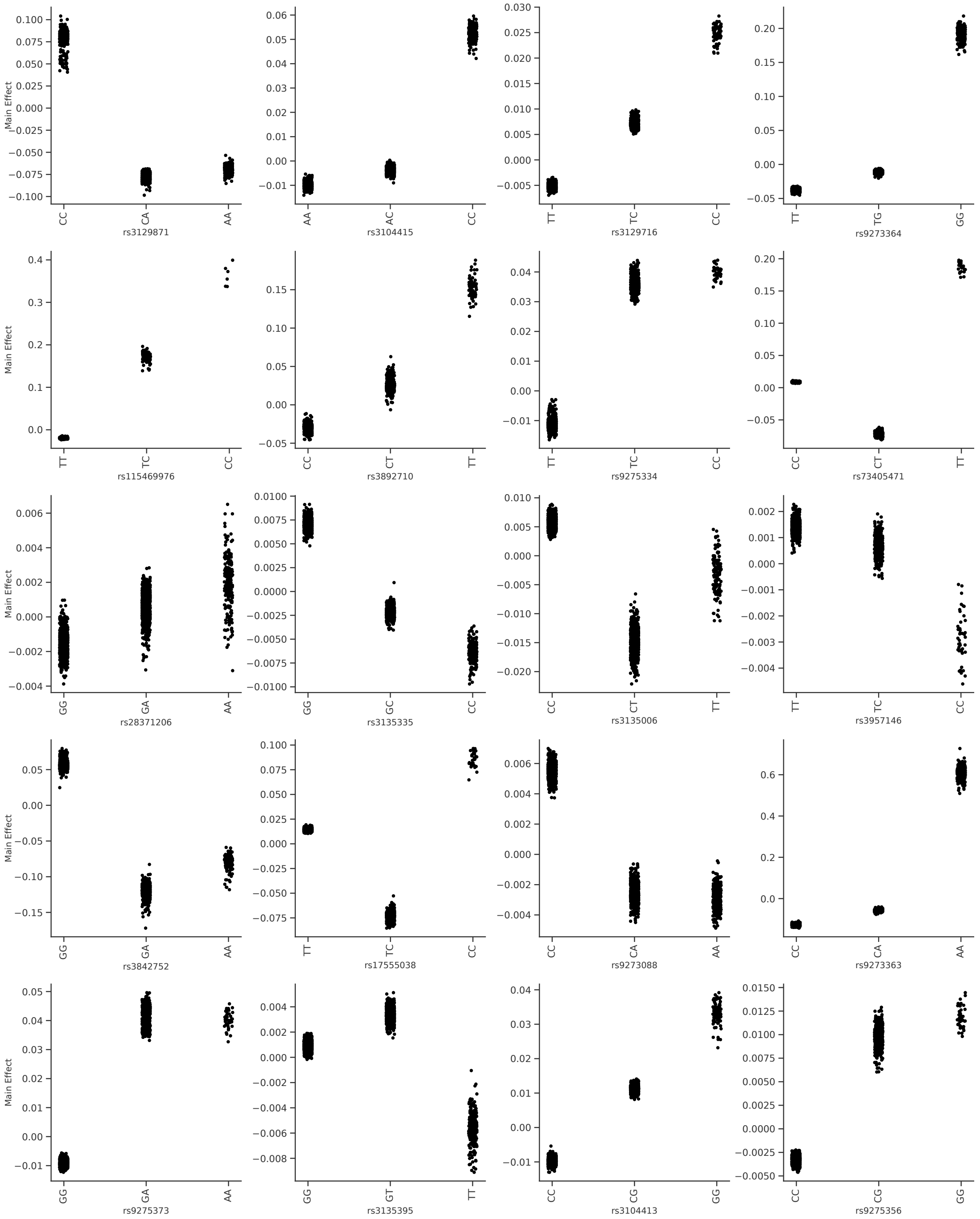


**Supplementary Figure 5.** Main effects of the top 20 most important identified from multiple GLN training runs for type 1 diabetes. Ten training runs with different seeds were performed with the GLN model, and the top 200 SNPs across all runs used as candidates for the analysis. A gradient boosted decision trees models was trained on the top SNPs and analyzed for both main and interaction effects. The SNPs show both additive effects also non-additive effects where the heterozygote and homozygote show opposite effects.

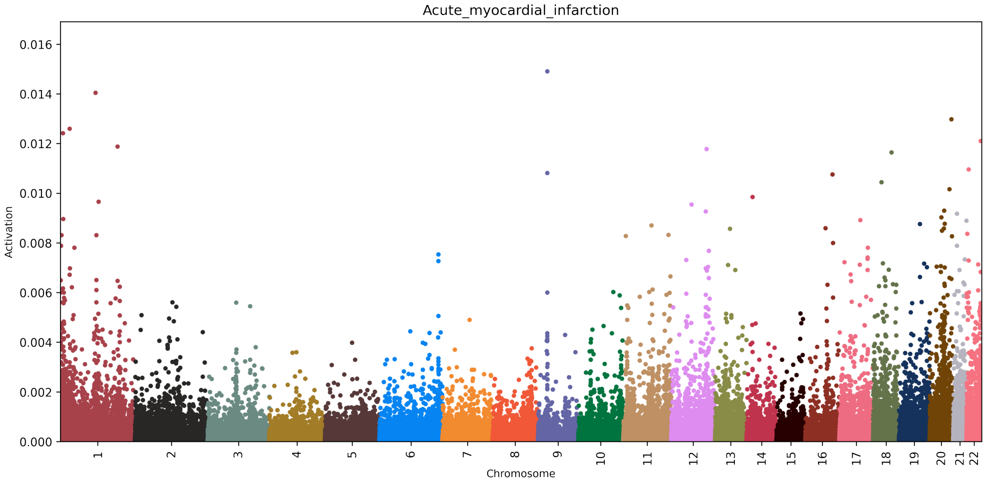


**Supplementary Figure 6.** SNP activation distribution using the GLN model for acute myocardial infarction. The values on the y-axis represent a given SNP's absolute influence on the model's raw output score (logit) for acute myocardial infarction.

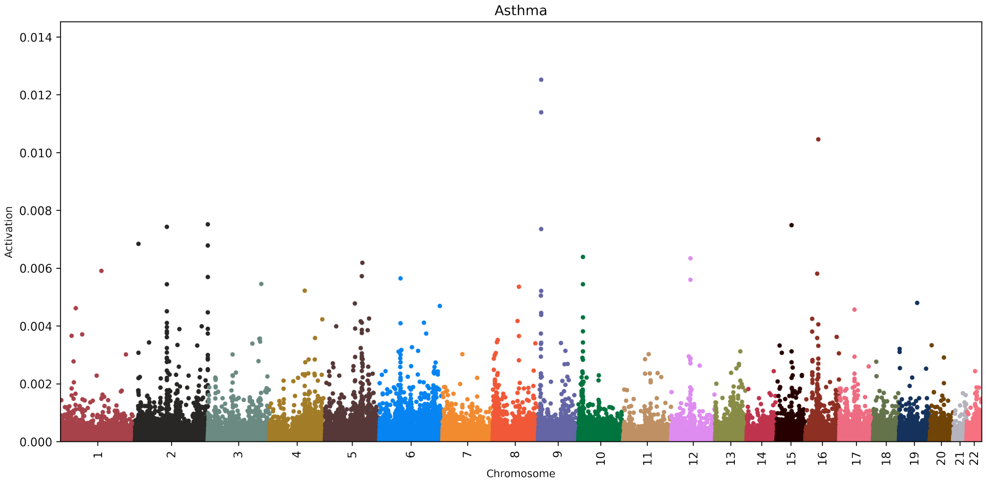


**Supplementary Figure 7.** SNP activation distribution using the GLN model for asthma. The values on the y-axis represent a given SNP's absolute influence on the model's raw output score (logit) for asthma.

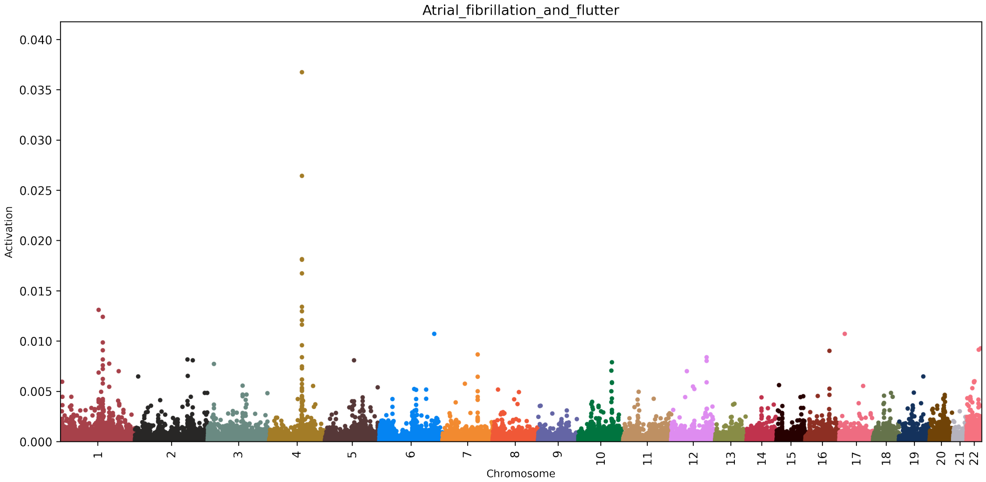


**Supplementary Figure 8.** SNP activation distribution using the GLN model for atrial fibrillation and flutter. The values on the y-axis represent a given SNP's absolute influence on the model's raw output score (logit) for atrial fibrillation and flutter.

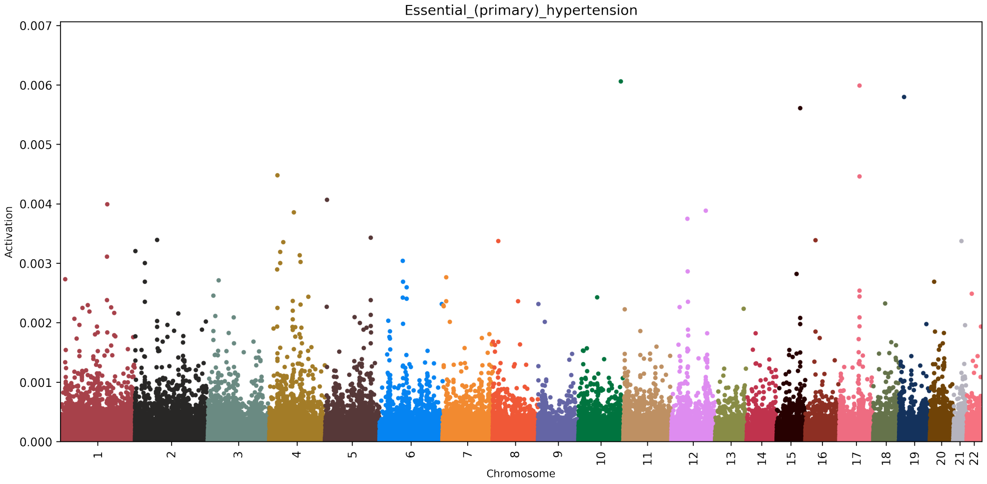


**Supplementary Figure 9.** SNP activation distribution using the GLN model for hypertension. The values on the y-axis represent a given SNP's absolute influence on the model's raw output score (logit) for hypertension.

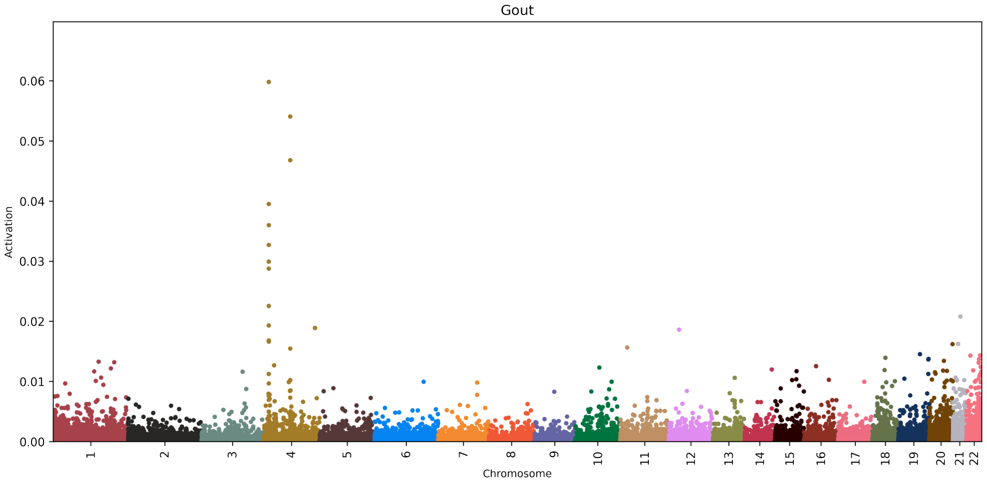


**Supplementary Figure 10.** SNP activation distribution using the GLN model for gout. The values on the y-axis represent a given SNP's absolute influence on the model's raw output score (logit) for gout.

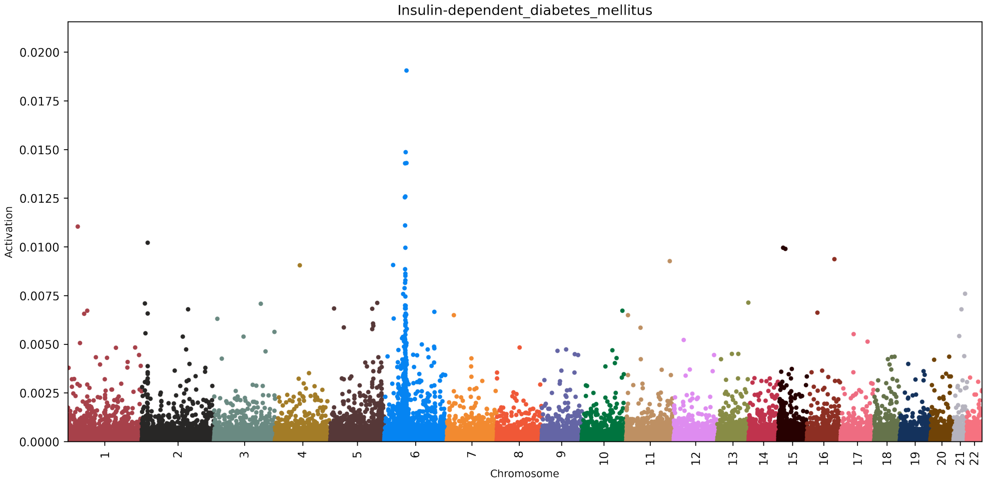


**Supplementary Figure 11.** SNP activation distribution using the GLN model for type 1 diabetes. The values on the y-axis represent a given SNP's absolute influence on the model's raw output score (logit) for Type 1 Diabetes.

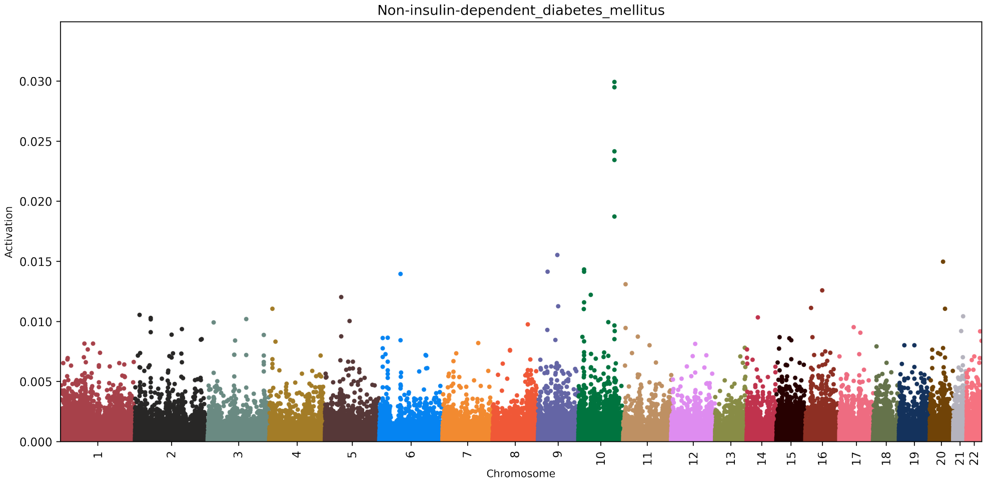


**Supplementary Figure 12.** SNP activation distribution using the GLN model for type 2 diabetes. The values on the y-axis represent a given SNP's absolute influence on the model's raw output score (logit) for Type 2 Diabetes.

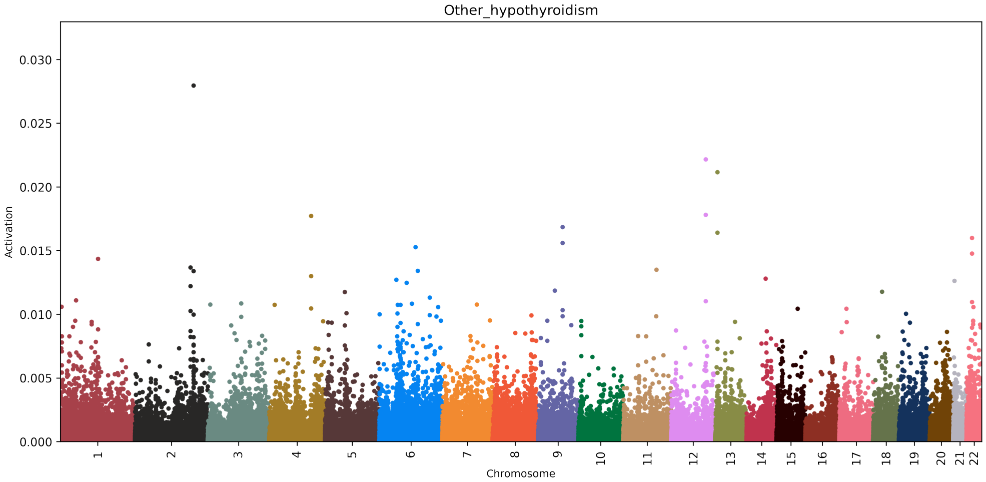


**Supplementary Figure 13.** SNP activation distribution using the GLN model for hypothyroidism. The values on the y-axis represent a given SNP's absolute influence on the model's raw output score (logit) for hypothyroidism.

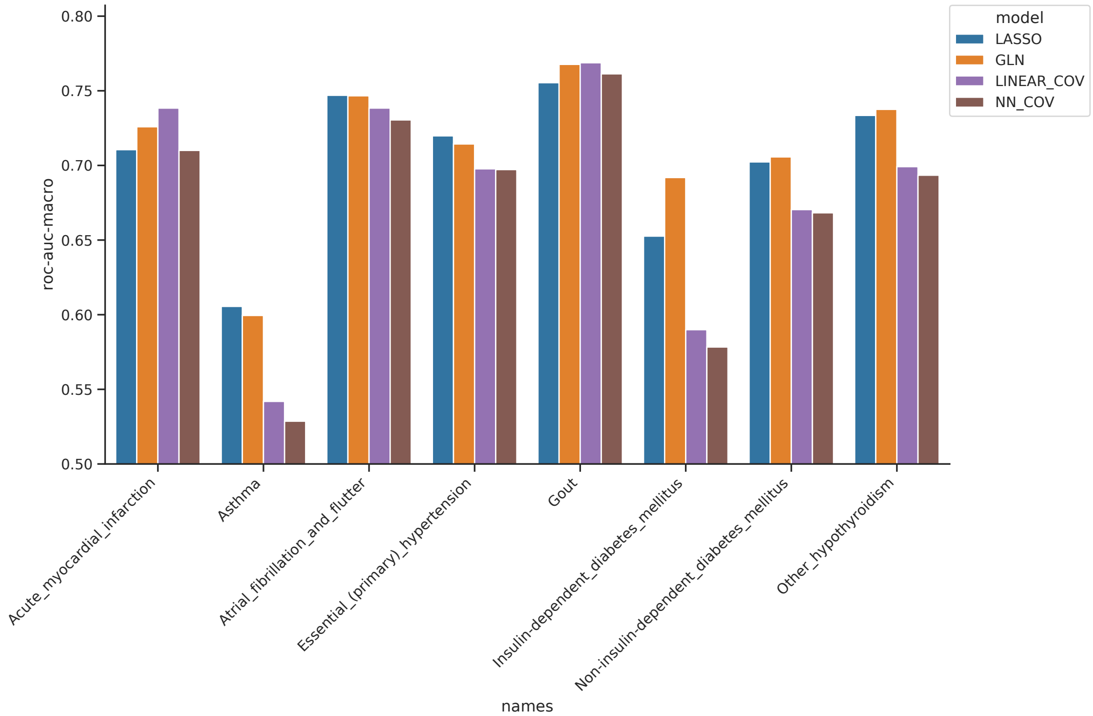


**Supplementary Figure 14.** Comparison of using the LASSO (blue) and GLN (orange) models with the covariates age, sex and first 10 PCs and genotype against only using the covariates modelled with a linear (purple) and neural network based model (brown), measured in ROC-AUC for the 8 benchmark traits. When only using the covariates, the same model architecture is used but with the genotype modality omitted.

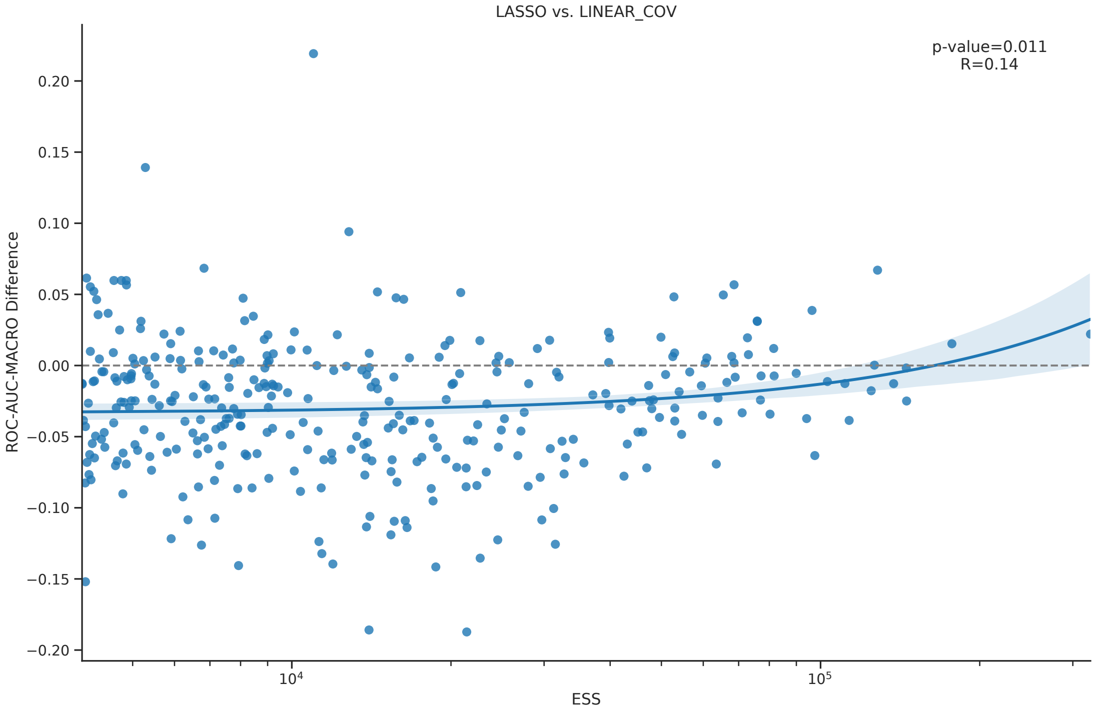


**Supplementary Figure 15.** Difference in performance of LASSO model versus linear covariate based model (using sex, age and the first 10 genotype principal components) as a function of effective sample size (ESS). A positive difference indicates that the LASSO model performed better on the test set compared to the linear covariate based model.

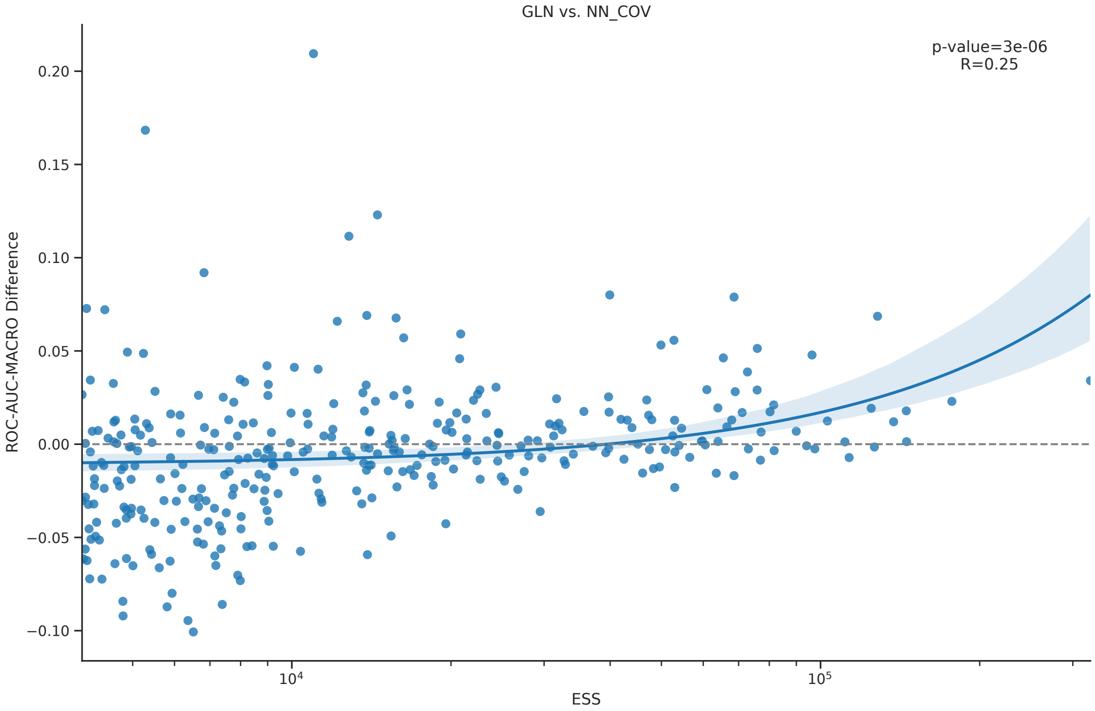


**Supplementary Figure 16.** Difference in performance of GLN model versus a neural network (NN) covariate based model (using sex, age and the first 10 genotype principal components) as a function of effective sample size (ESS). A positive difference indicates that the GLN model performed better on the test set compared to the NN covariate based model.

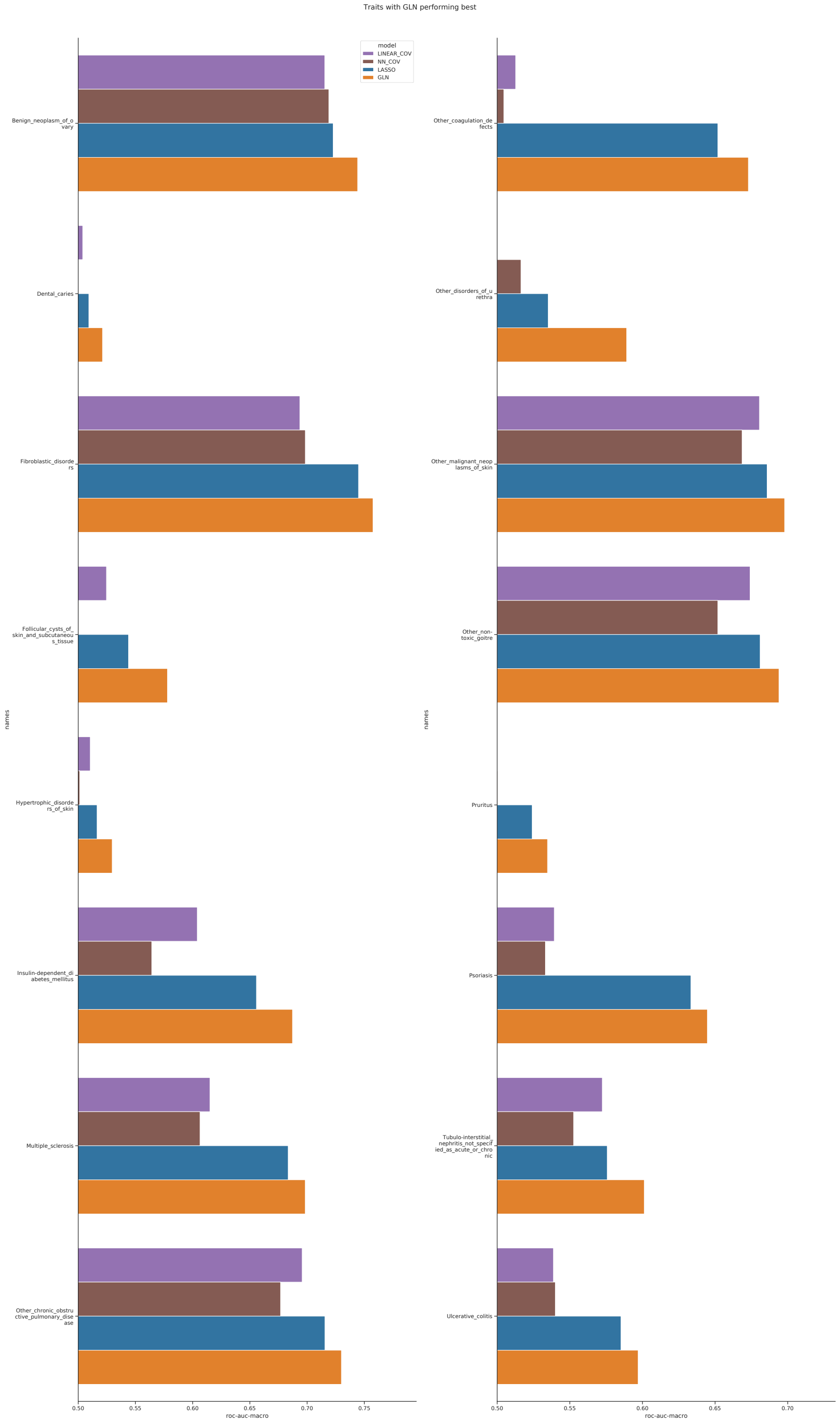


**Supplementary Figure 17.** Traits in the large scale training where the GLN (orange) model showed the best performance (with a ROC-AUC difference greater than 0.01) and both GLN and LASSO (blue) had a better performance than either of the linear (purple) or neural network (brown) base covariates models on the held-out test set. All models were adjusted for sex, age and the first 10 principal components.

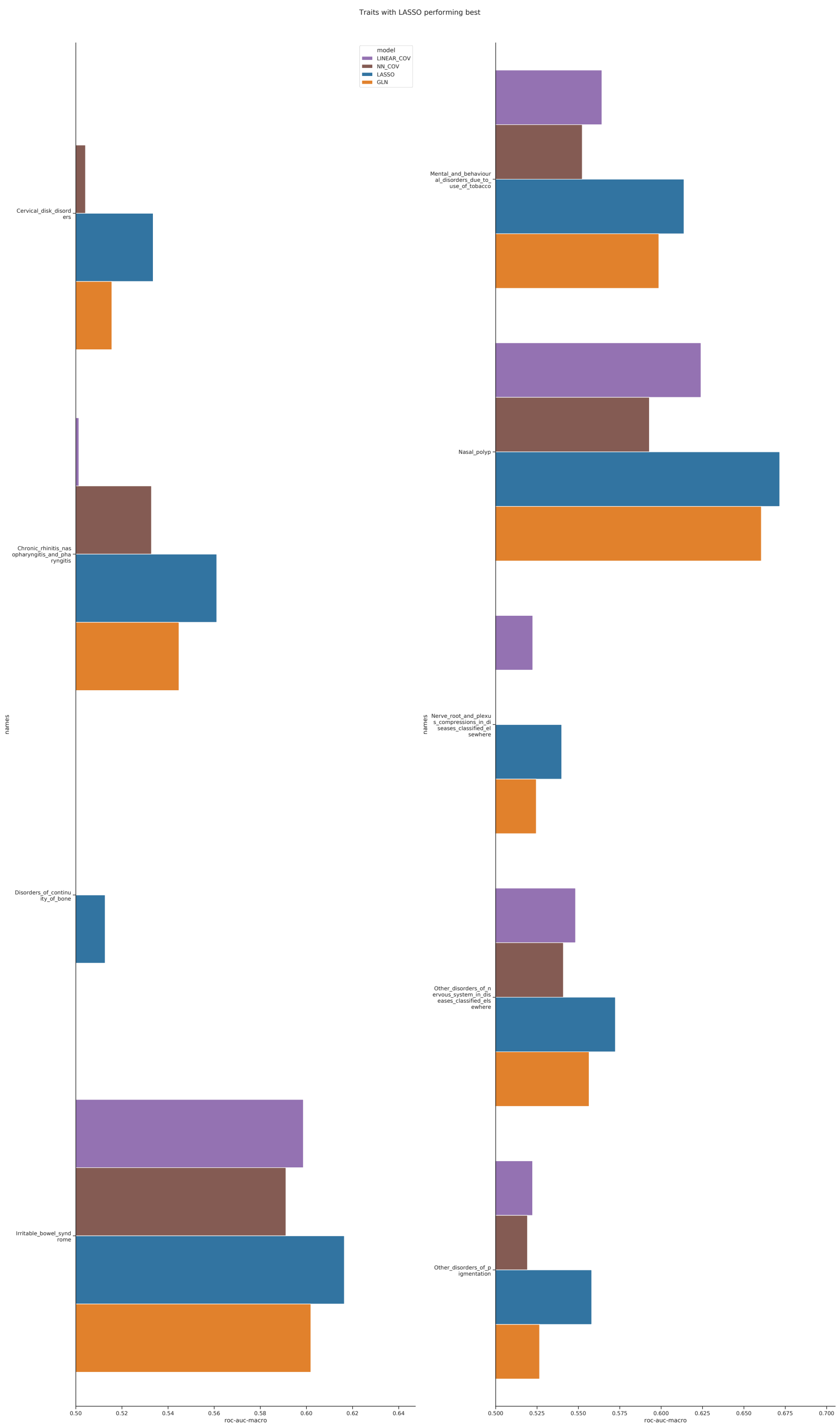


**Supplementary Figure 18.** Traits in the large scale training where the LASSO (blue) model showed the best performance (with a ROC-AUC difference greater than 0.01) and both GLN (orange) and LASSO had a better performance than either of the linear (purple) or neural network (brown) base covariates models on the test set. All models were adjusted for sex, age and the first 10 principal components.

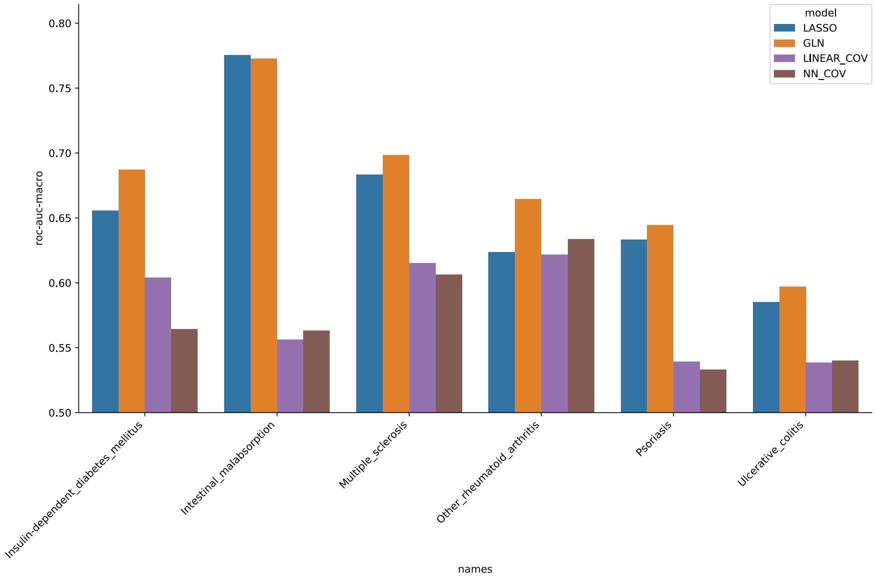


**Supplementary Figure 19.** Comparison of using the LASSO (blue) and GLN (orange) models with the covariates age, sex and first 10 PCs and genotype against only using the covariates modelled with a linear (purple) and neural network based model (brown), for autoimmune traits previously researched for interaction effects. When only using the covariates, the same model architecture is used but with the genotype modality omitted. Performance is measured in ROC-AUC on the held-out test set.

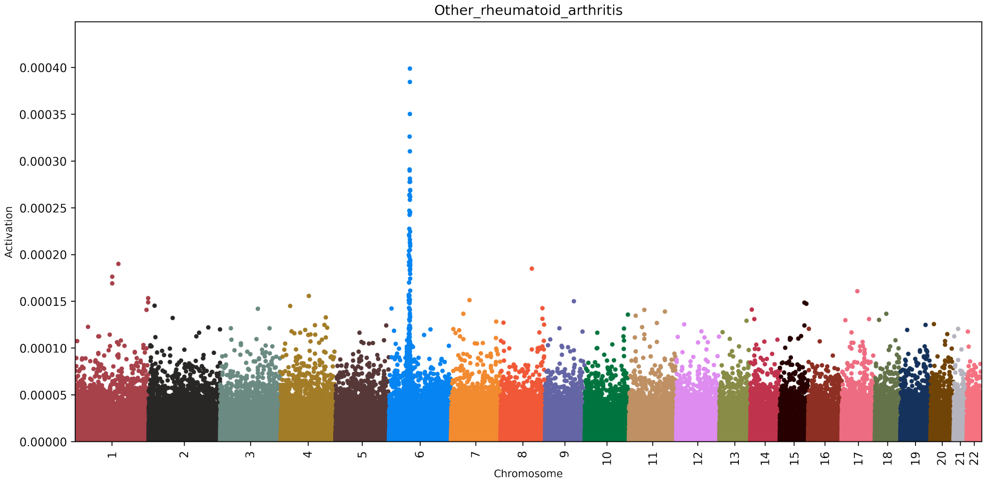


**Supplementary Figure 20.** SNP activation distribution using the LASSO model for rheumatoid arthritis. The values on the y-axis represent a given SNP's absolute influence on the model's raw output score (logit) for rheumatoid arthritis.

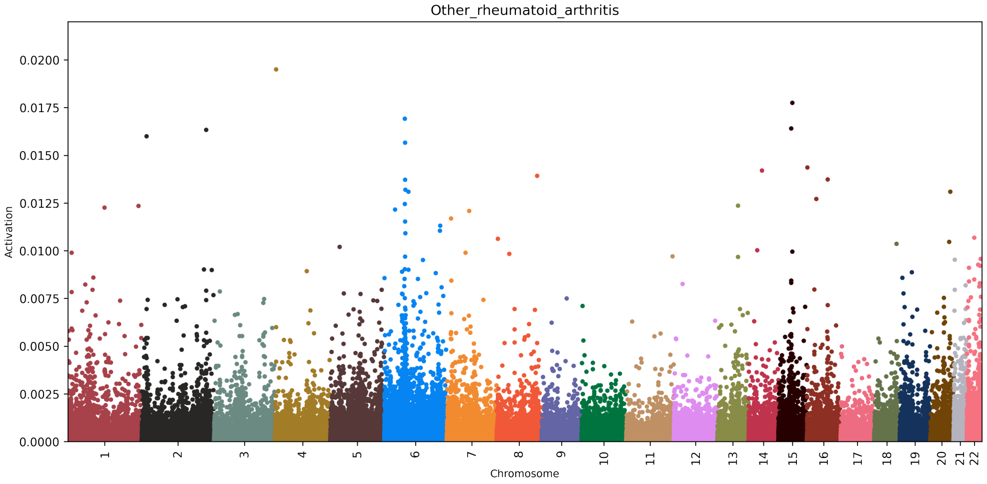


**Supplementary Figure 21.** SNP activation distribution using the GLN model for rheumatoid arthritis. The values on the y-axis represent a given SNP's absolute influence on the model's raw output score (logit) for rheumatoid arthritis.

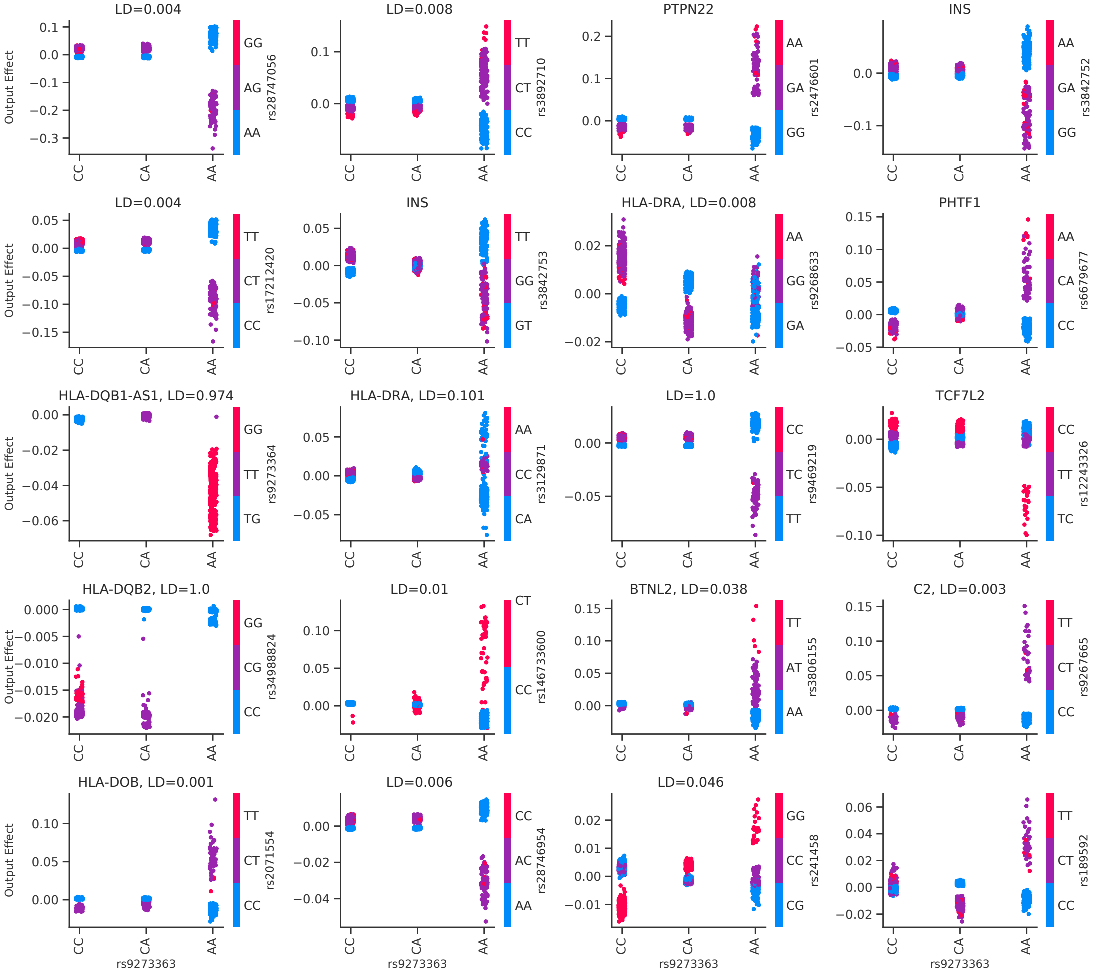


**Supplementary Figure 22.** Interaction effects between rs9273363 and the other top 20 most important SNPs identified when training the gradient boosted decision trees on the top 200 SNPs from the GLN model for T1D. The x-axis corresponds to the rs9273363 alleles, while the y-axis corresponds to the effect of the interaction on the raw gradient boosted decision trees (GBDT) model output (logit). Each dot in the figure represents one sample, and the dot color indicates the allele of the SNP interacting with rs9273363. If the SNP interacting with rs9273363 was mapped to a gene, it is shown above the relevant sub-figure. Additionally, if the interacting SNP resides on chr6 as rs9273363 does, the linkage disequilibrium (LD) value was computed (in $R^2$) based on the British in England and Scotland population of LDLink[100] is shown.

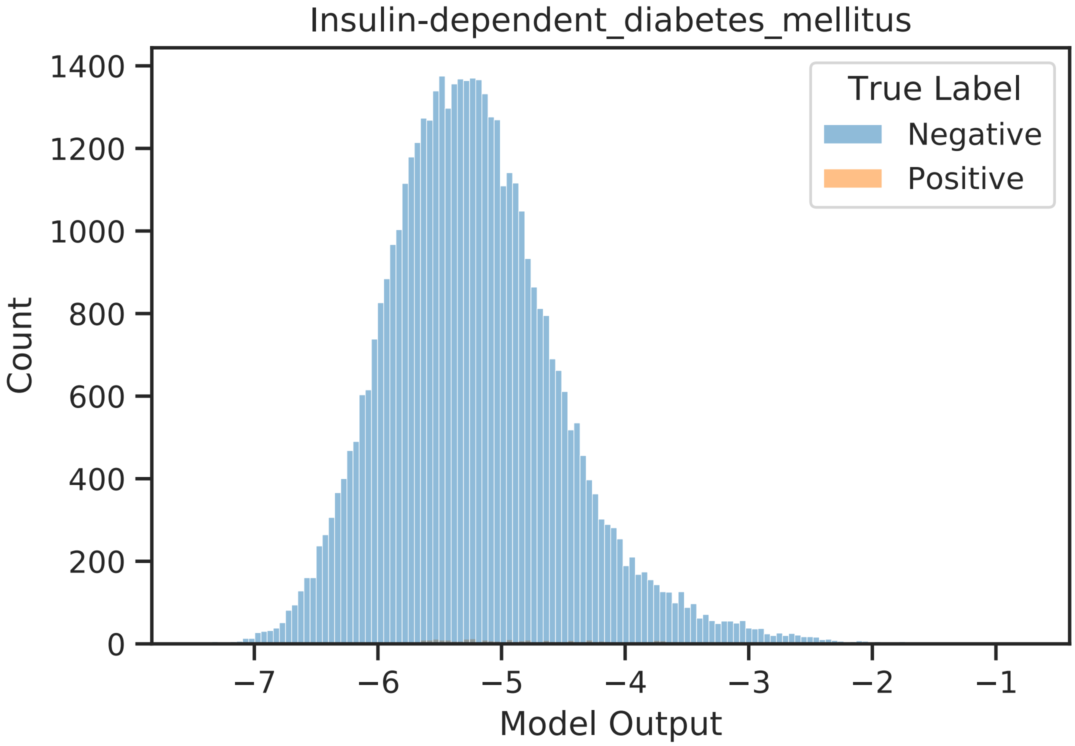


**Supplementary Figure 23.** Output distribution of the gradient boosted decision trees (GBDT) model trained on the top 200 GLN activated SNPs, where the top 200 SNPs are chosen from the average activation of 10 GLN training runs with different seeds each.

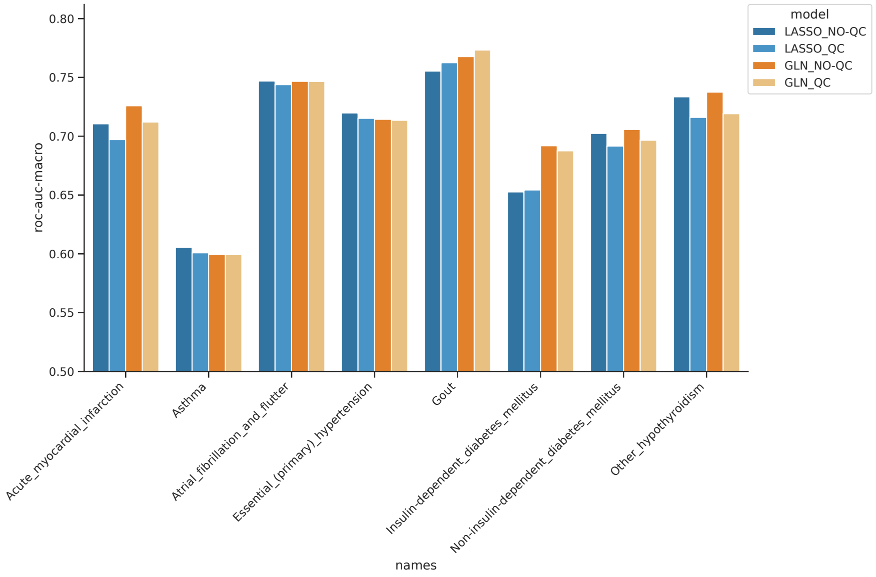


**Supplementary Figure 24.** Comparison of using traditional genotype pre-processing (QC) or not (NO-QC) (Methods) for 8 benchmark traits using the LASSO model trained with NO-QC data (blue), LASSO trained with QC data (light blue), GLN trained with NO-QC data (orange) and GLN trained with QC data (light orange).

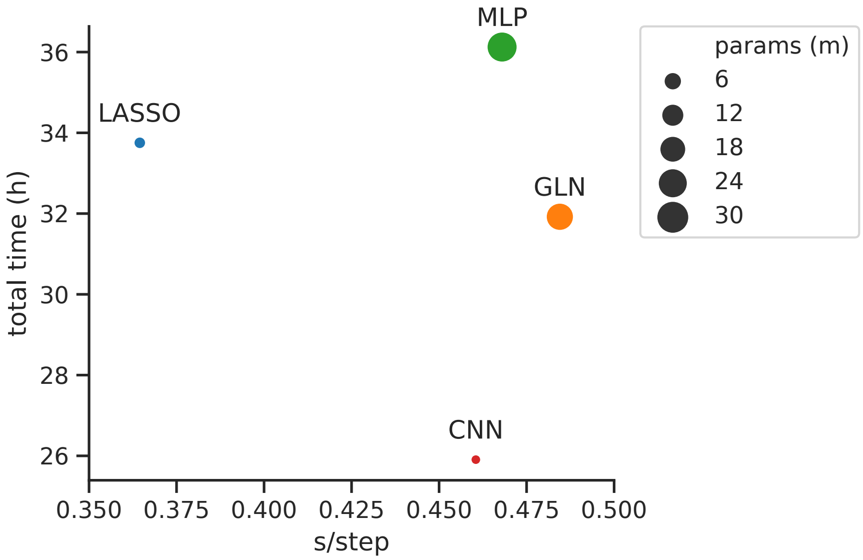


**Supplementary Figure 25.** Comparison of model total training time, training latency (seconds per batch of 64 samples) and number of parameters for the LASSO (blue), GLN (orange), MLP (green) and CNN (red) models. The y-axis represents total training time for all 8 benchmark traits, and the x-axis the time it took for a given model to do one forward and backwards pass of 64 samples. The point size is according to the number of model parameters.

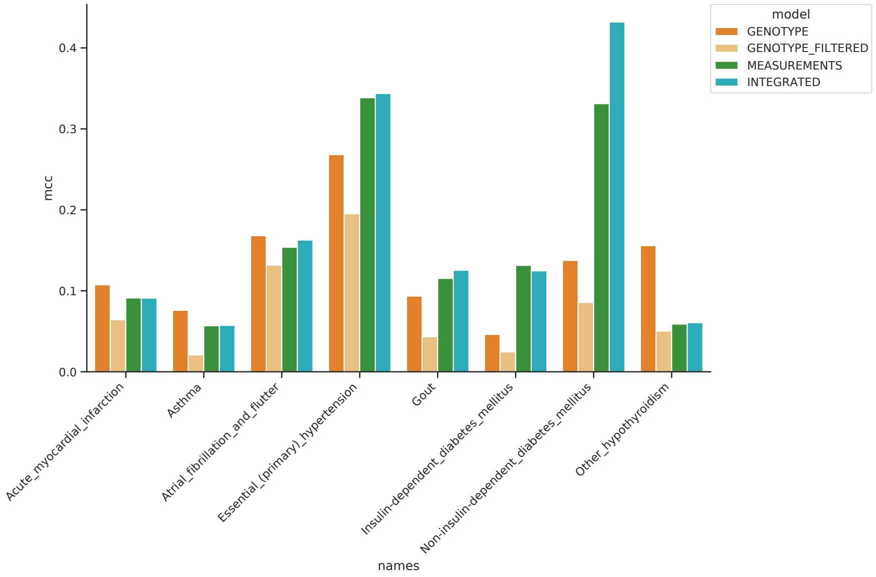


**Supplementary Figure 26.** Comparison of model performance using Genotype (orange), Genotype Filtered (light orange), Measurement (green) and Integrated (teal) data in MCC on the held-out test set.

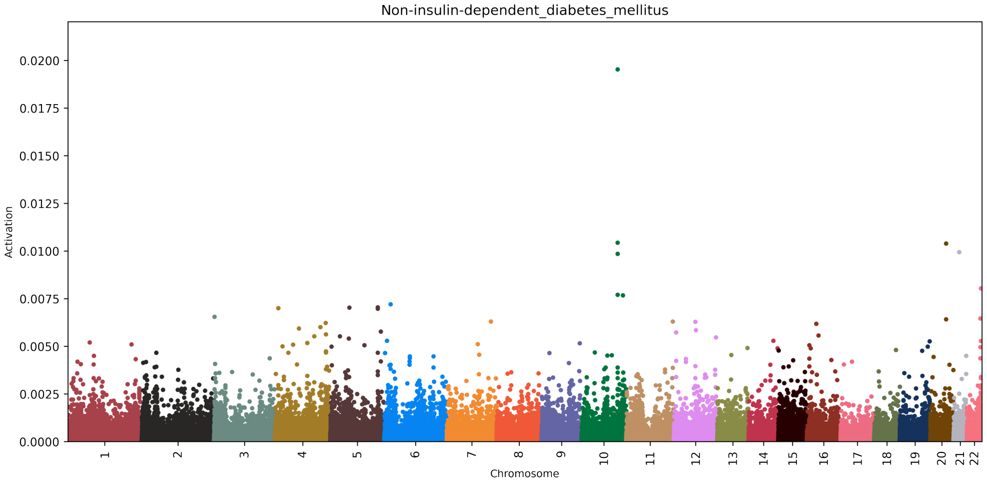


**Supplementary Figure 27.** SNP activation distribution using the GLN model for type 2 diabetes, when including clinical and biochemical measurements. The values on the y-axis represent a given SNP's absolute influence on the model's raw output score (logit) for type 2 diabetes.

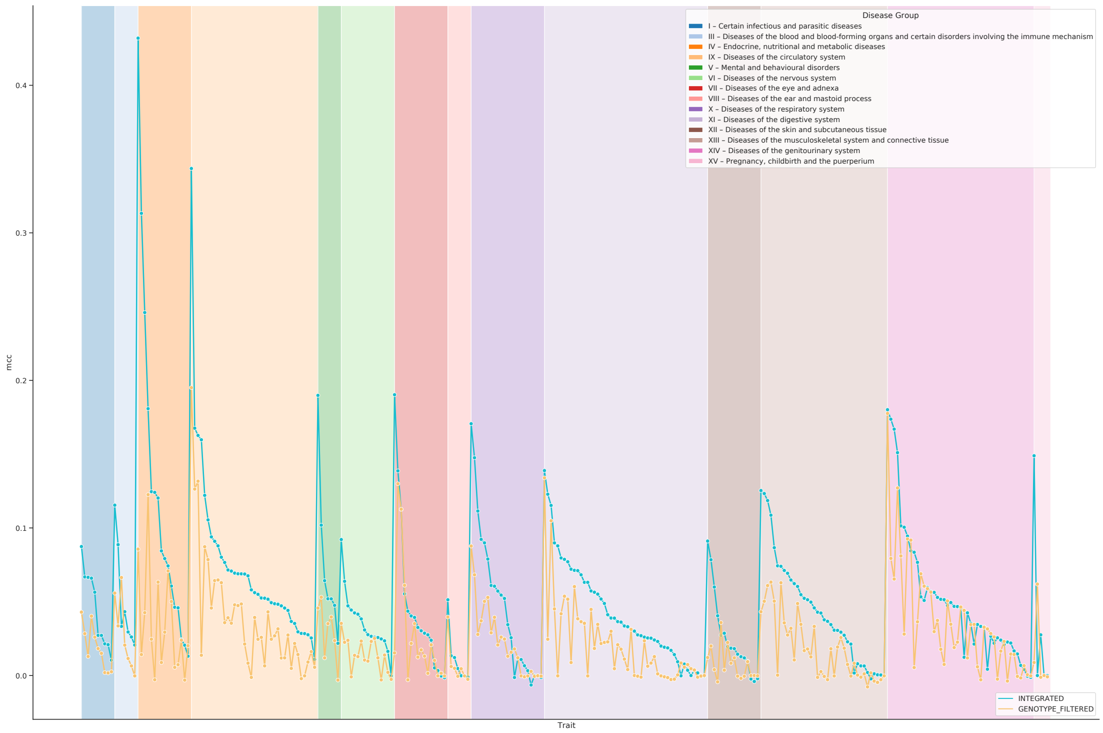


**Supplementary Figure 28.** Summary of MCC performance on the held-out test set across all the 290 traits that had a time measured column associated with them, with Integrated data (teal) compared with Genotype Filtered data (light orange), filtered for time of diagnosis. The different background colors represent different ICD-10 chapters.

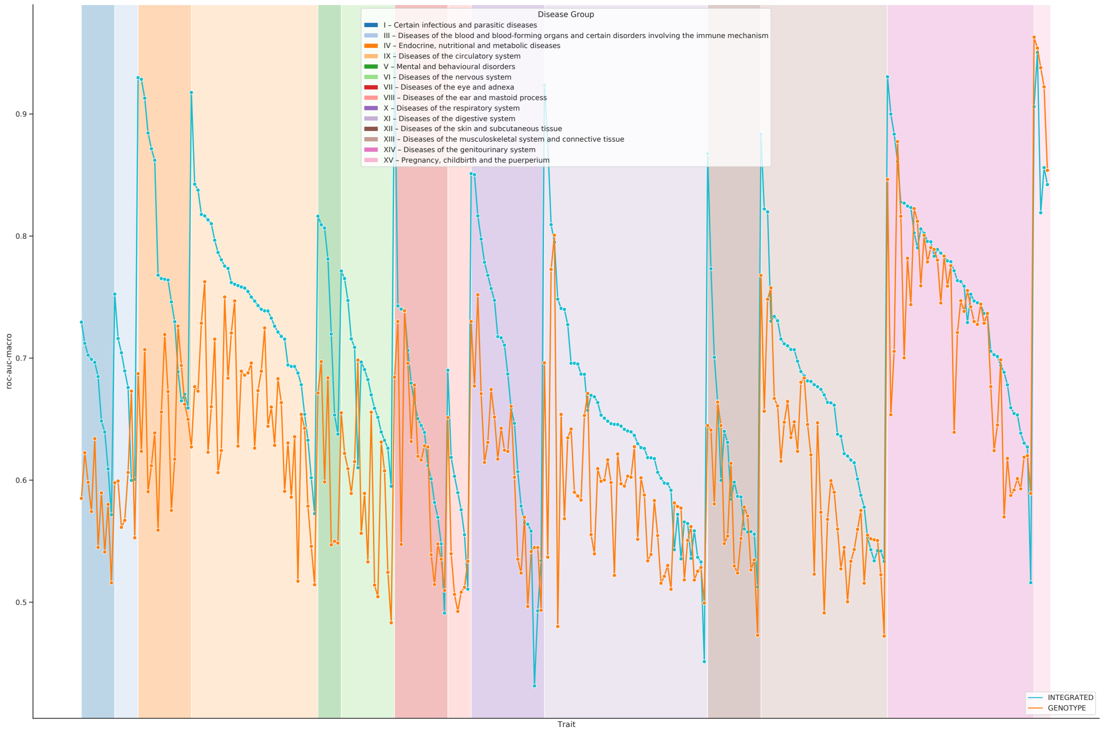


**Supplementary Figure 29.** Summary of ROC-AUC performance on the held-out test set across all the 290 traits that had a time measured column associated with them, with Integrated data (teal) compared with Genotype data (orange), filtered for time of diagnosis. The different background colors represent different ICD-10 chapters.

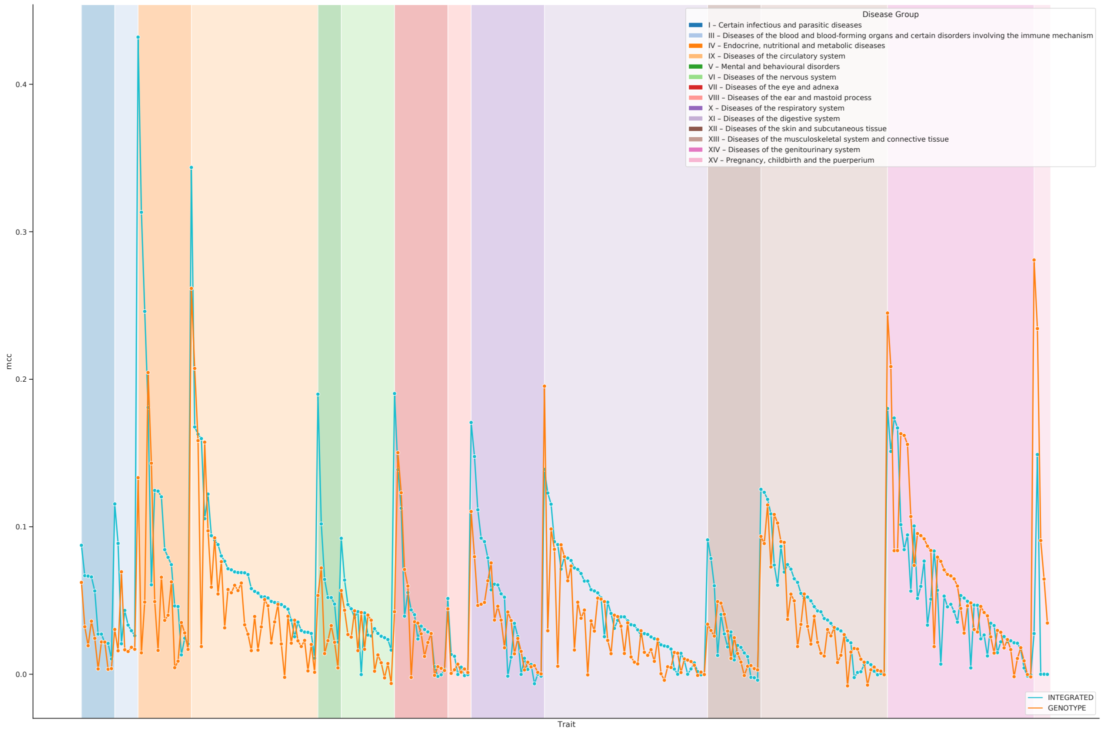


**Supplementary Figure 30.** Summary of MCC performance on the held-out test set across all the 290 traits that had a time measured column associated with them, with Integrated data (teal) compared with Genotype data (orange), filtered for time of diagnosis. The different background colors represent different ICD-10 chapters.

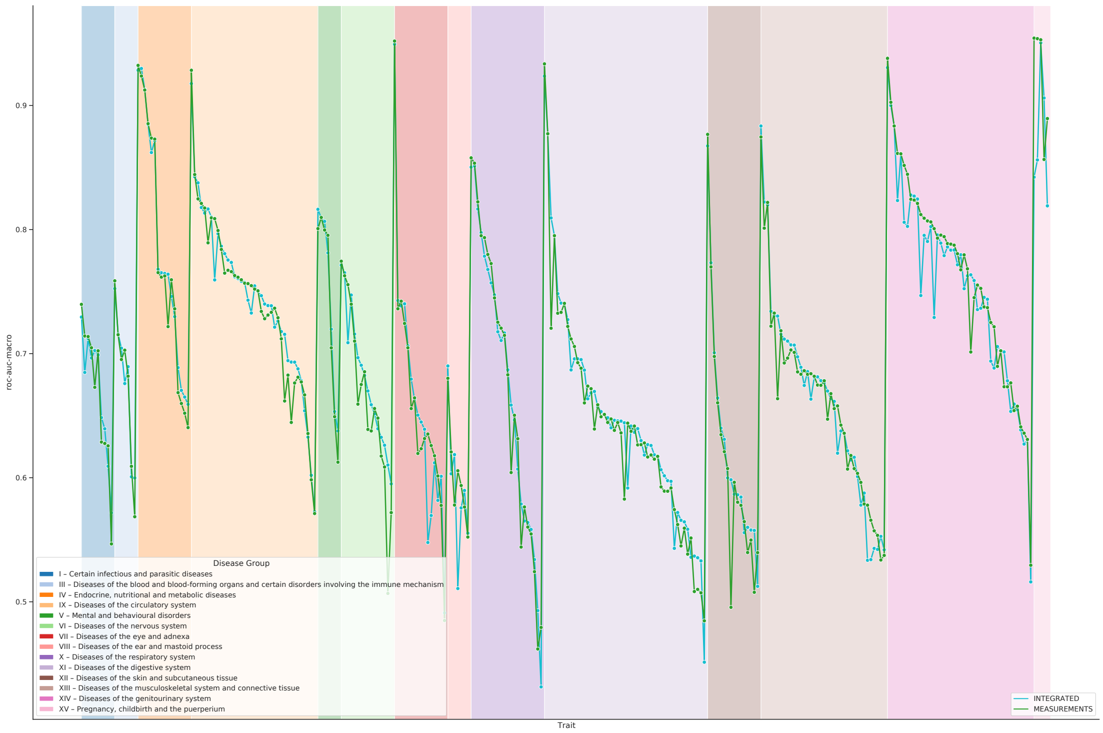


**Supplementary Figure 31.** Summary of ROC-AUC performance on the held-out test set across all the 290 traits that had a time measured column associated with them, with Integrated data (teal) compared with Measurement data (green), filtered for time of diagnosis. The different background colors represent different ICD-10 chapters.

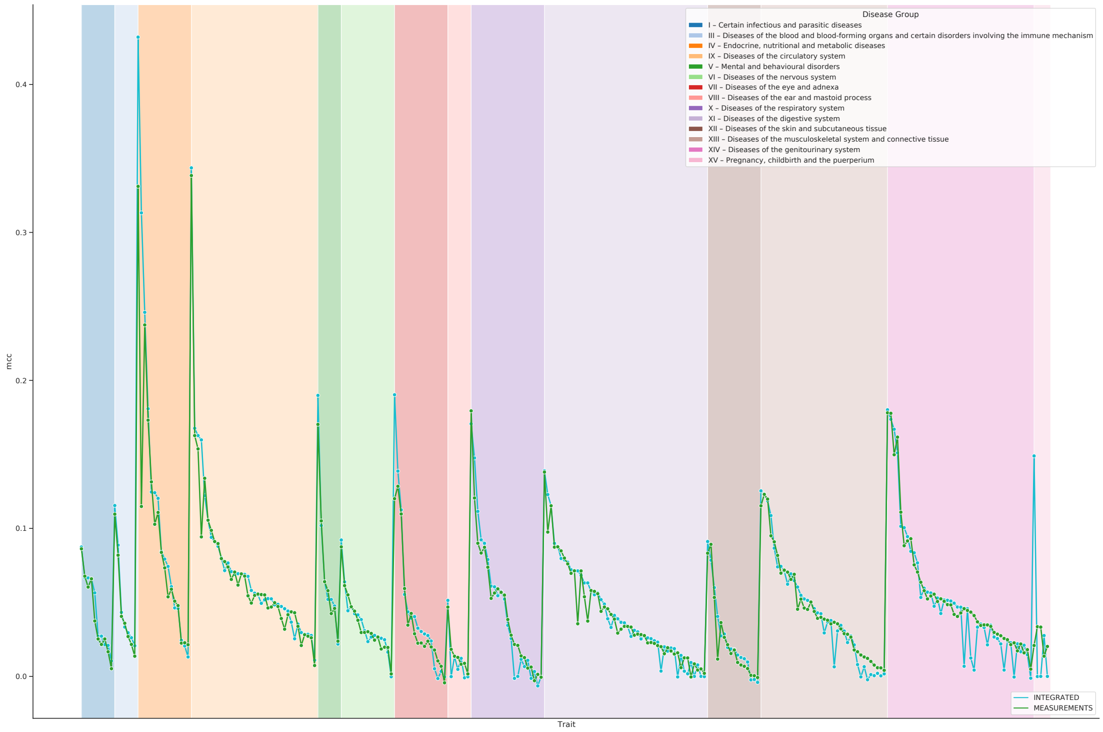


**Supplementary Figure 32.** Summary of MCC performance on the held-out test set across all the 290 traits that had a time measured column associated with them, with Integrated data (teal) compared with Measurement data (green), filtered for time of diagnosis. The different background colors represent different ICD-10 chapters.

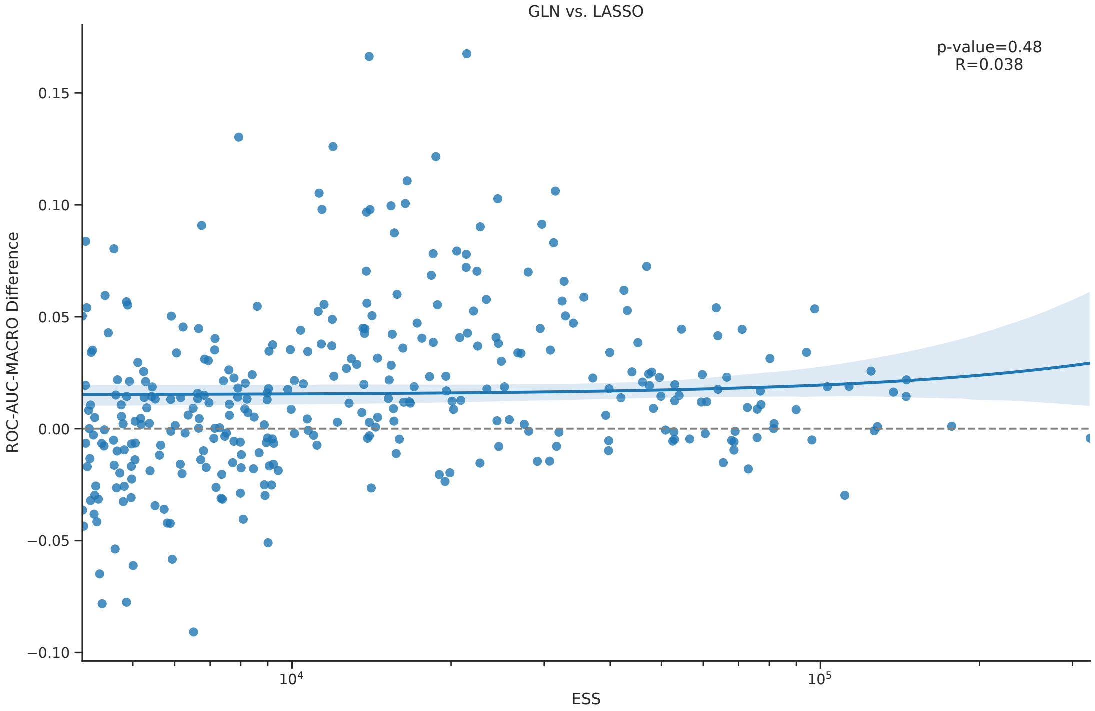


**Supplementary Figure 33.** Difference in performance of GLN model versus the LASSO model as a function of effective sample size (ESS). A positive difference indicates that the GLN model performed better on the test set compared to the LASSO.

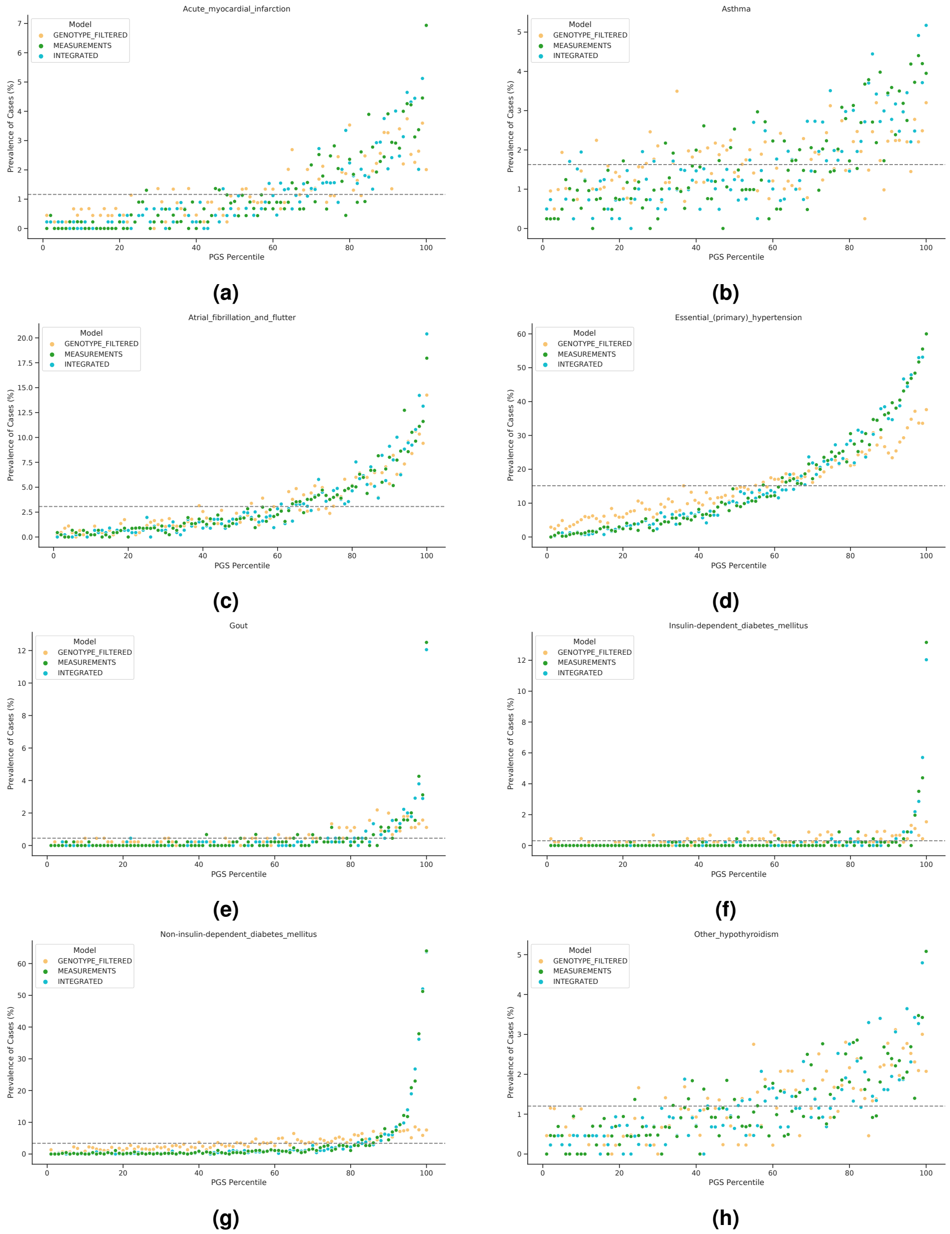


**Supplementary Figure 34.** Prevalence plots comparing models using Genotype Filtered (light orange), Measurement (green) and Integrated (teal) data, using the GLN model. The dashed line represents the average prevalence in the test set, after filtering for time of diagnosis.

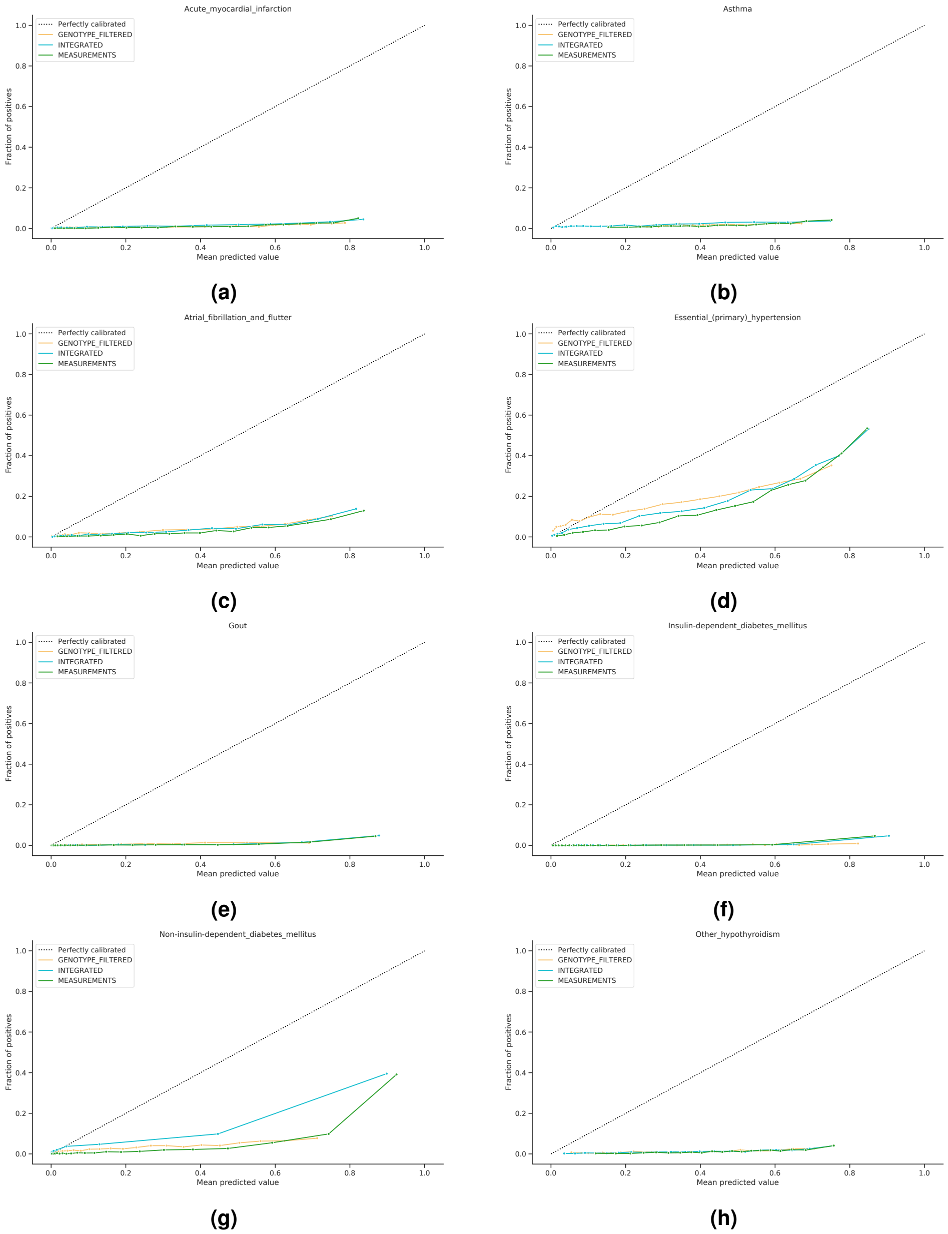


**Supplementary Figure 35.** Calibration plots comparing Genotype filtered (light orange), Measurements (green) and Integrated (teal) input data, using the GLN model.

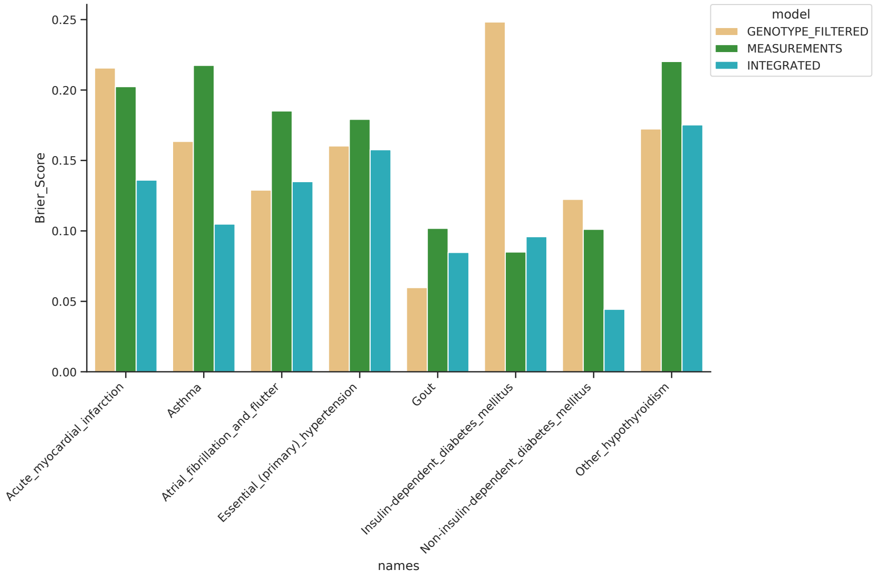


**Supplementary Figure 36.** Comparison of Genotype filtered (light orange), Measurements (green) and Integrated (teal) input data using Brier Score (lower better), using the GLN model.

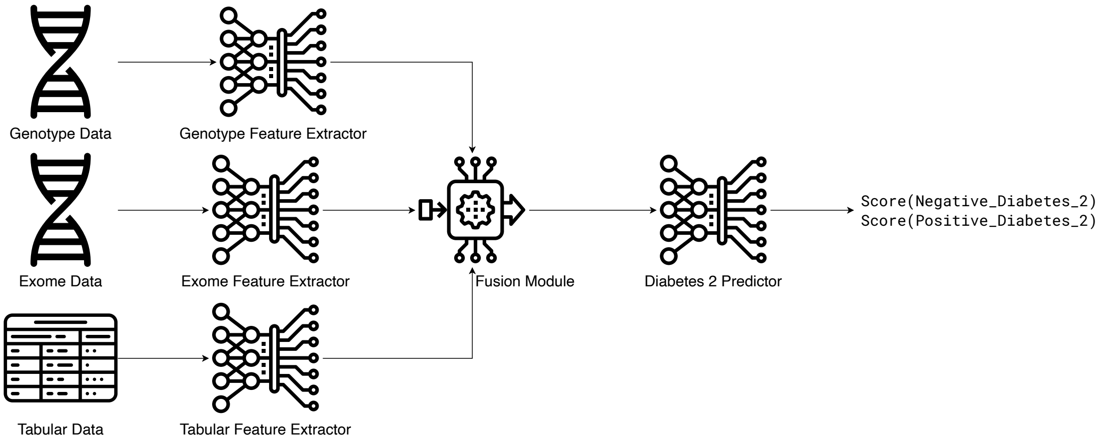


**Supplementary Figure 37.** Example overall architecture. Different sources and/or modalities can go through different NN feature extractors, which generate different intermediate representation for each input source. The representations get fused in a fusion module. The fused representation finally goes through a NN classifier, which generates the final prediction output.

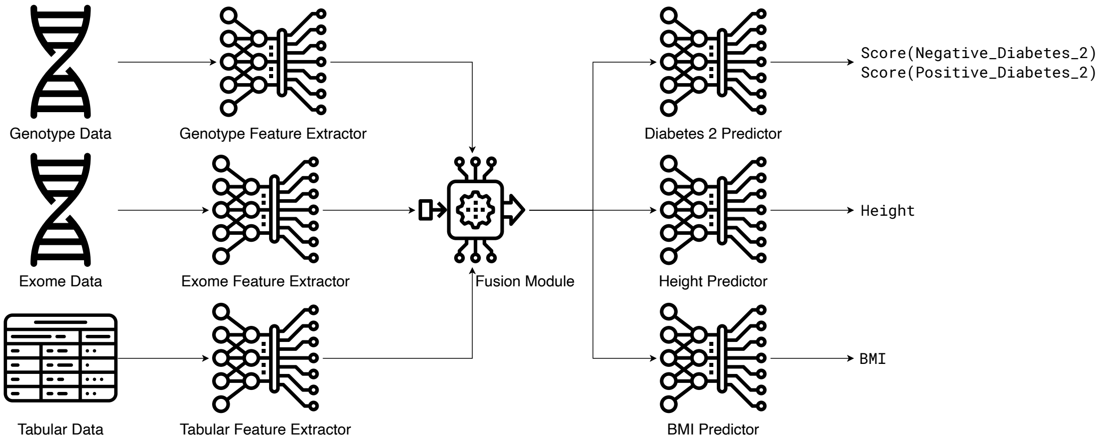


**Supplementary Figure 38.** Example overall architecture for multi-task learning. The figure is analogous to Supplementary Figure 37, but instead of predicting one output variable from the fused representation, it is passed to multiple predictors.

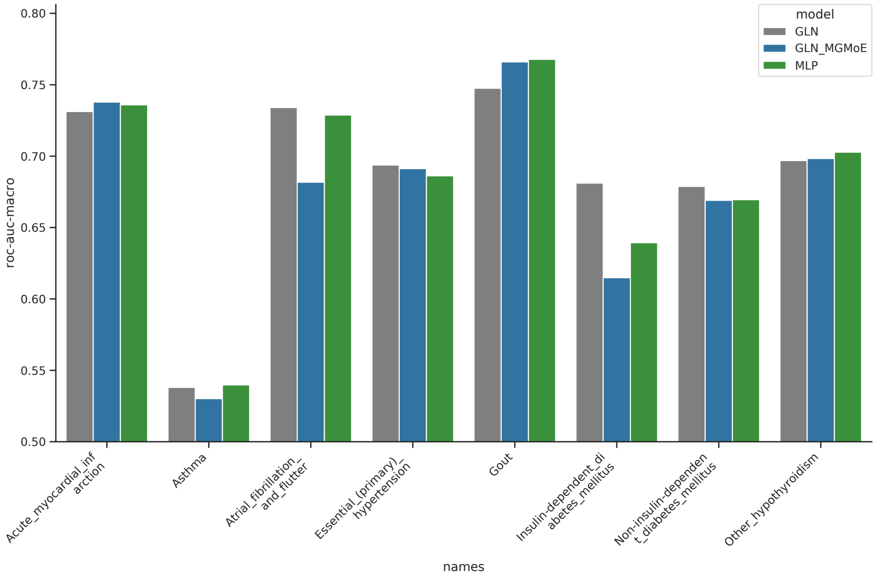


**Supplementary Figure 39.** Comparison of different MT architectures on the 8 benchmark traits. The GLN (gray), GLN_MGMoE (blue) and MLP (green) models had an average ROC-AUC of 0.688, 0.674 and 0.684 respectively.

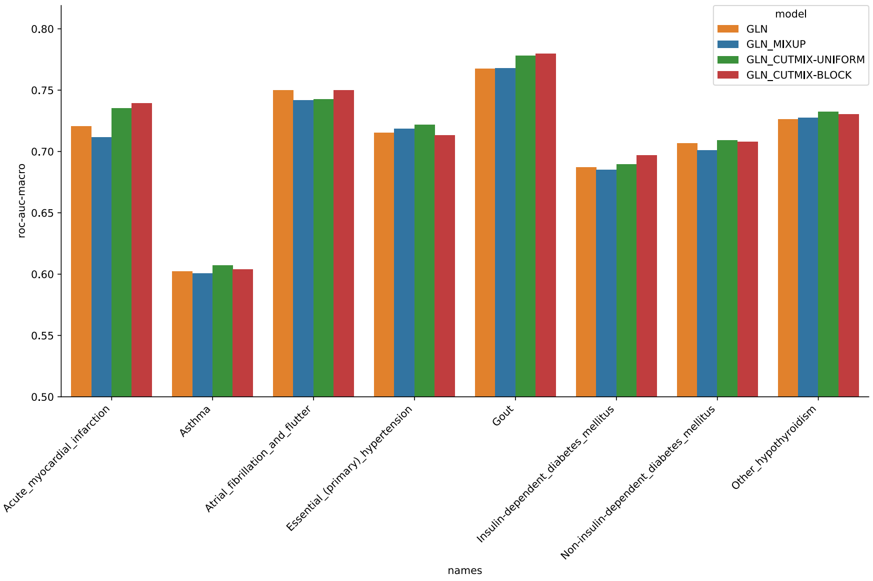


**Supplementary Figure 40.** Comparison of different augmentation strategies implemented by the framework. While Mixup (blue) mixes inputs with linear interpolations, CutMix does so in a cut-and-paste manner. The difference between CutMix-Uniform (green) and Cutmix-Block (red) is they mix a set of uniformly distributed single SNPs and a single block of the genotype data, respectively. The performance of the GLN model without any mixing augmentations (orange) is shown for comparison.

| Simulated Data Type | Model | Validation $R^2$ | Parameters |
|---|---|---|---|
| Additive | Linear | 0.9999 | 4001 |
| Additive | CNN | 0.9977 | 20877 |
| Additive | MLP | 0.9988 | 65732 |
| Additive | GLN | 0.9978 | 12810 |
| Mix | Linear | 0.7517 | 4001 |
| Mix | CNN | 0.9779 | 20877 |
| Mix | MLP | 0.9787 | 65732 |
| Mix | GLN | 0.9792 | 12810 |
| XOR | Linear | -0.0291 | 4001 |
| XOR | CNN | 0.9611 | 20877 |
| XOR | MLP | 0.9479 | 65732 |
| XOR | GLN | 0.958 | 12810 |

**Supplementary Table 1.** Model comparison using simulated data. Mix refers to mixed effects of additive and XOR simulated SNP interactions. 12000 samples with 1000 SNPs each were simulated.

| L1 | LR | NA | Validation ROC-AUC | Test ROC-AUC |
|---|---|---|---|---|
| 1e-04 | 5e-06 | 0.0 | 0.6638 | |
| 1e-04 | 5e-05 | 0.0 | 0.6617 | |
| 1e-04 | 5e-04 | 0.0 | 0.6604 | |
| 1e-04 | 5e-03 | 0.0 | 0.5058 | |
| 1e-03 | 5e-06 | 0.0 | 0.6707 | |
| 1e-03 | 5e-05 | 0.0 | 0.6685 | |
| 1e-03 | 5e-04 | 0.0 | 0.6404 | |
| 1e-03 | 5e-03 | 0.0 | 0.5057 | |
| 1e-02 | 5e-06 | 0.0 | 0.6511 | |
| 1e-02 | 5e-05 | 0.0 | 0.6535 | |
| 1e-02 | 5e-04 | 0.0 | 0.6319 | |
| 1e-02 | 5e-03 | 0.0 | 0.5974 | |
| 1e-01 | 5e-06 | 0.0 | 0.5907 | |
| 1e-01 | 5e-05 | 0.0 | 0.5811 | |
| 1e-01 | 5e-04 | 0.0 | 0.5544 | |
| 1e-01 | 5e-03 | 0.0 | 0.534 | |
| 1e-03 | 5e-05 | 0.1 | 0.6742 | |
| 1e-03 | 5e-05 | 0.2 | 0.6736 | |
| 1e-03 | 5e-05 | 0.4 | 0.6765 | 0.6596 |

**Supplementary Table 2.** Comparison of various LASSO hyperparameter combinations tested when modelling on T1D. To reduce the chance of overfitting on the test set, only the model with the best performance on the validation set was considered, and its performance measured on the test set.

| rsID | Chr. | Gene | Top 3 LitVar | Top 3 DisGeNET |
|---|---|---|---|---|
| rs115469976 | 6 | | | |
| rs3892710 | 6 | | Inflammation, Zellweger Syndrome, Coronary Artery Disease | |
| rs2281390 | 6 | | | |
| rs58495148 | 6 | HLA-DRB6 | | Child Development Disorders, Pervasive, Lymphoma, Follicular, Multiple Sclerosis |
| rs73405471 | 6 | HLA-DQA1 | | Esophageal Achalasia, Celiac Disease, Diabetes Mellitus, Insulin-Dependent |
| rs984778 | 6 | | | |
| rs146733600 | 6 | | | |
| Affx-7384398 | 1 | | | |
| rs7583093 | 2 | | | |
| rs16954314 | 15 | LOC105370766 | | |
| Affx-37072118 | 6 | | | |
| rs79213347 | 15 | SNAP23 | | Liver Cirrhosis, Experimental, Myocardial Ischemia, Diabetes Mellitus, Non-Insulin-Dependent |
| rs72811854 | 16 | | | |
| rs2298483 | 11 | CHEK1 | | Malignant Neoplasm Of Breast, Breast Carcinoma, Mammary Neoplasms |
| rs196023 | 6 | CASC15 | | Neuroblastoma, Astigmatism, Regular Astigmatism - Corneal |
| rs4693440 | 4 | | | |
| rs9270656 | 6 | | | |
| rs9273364 | 6 | HLA-DQB1-AS1 | | Chronic Obstructive Airway Disease |
| rs35915063 | 6 | HLA-DQB1 | | Narcolepsy, Narcolepsy-Cataplexy Syndrome, Esophageal Achalasia |
| rs9273363 | 6 | HLA-DQB1-AS1 | Diabetes Mellitus, Arthritis, Rheumatoid, Autoimmune Diseases | Chronic Obstructive Airway Disease |

**Supplementary Table 3.** Cross-reference of the 20 most highly activated SNPs and known variant and gene associations for the GLN model trained on type 1 diabetes. The top 3 LitVar values were selected according to the most co-occuring disease entities. For DisGeNET, the top three scoring gene-disease DisGeNET association scores were used.

| rsID | Chr. | Gene | Top 3 LitVar | Top 3 DisGeNET |
|---|---|---|---|---|
| rs3129871 | 6 | HLA-DRA | Parkinson Disease, Multiple Sclerosis, Supranuclear Palsy, Progressive | Parkinson Disease, Multiple Sclerosis, Alcoholic Intoxication, Chronic |
| rs3104415 | 6 | | | |
| rs3129716 | 6 | | Retinoschisis | |
| rs9273364 | 6 | HLA-DQB1-AS1 | | Chronic Obstructive Airway Disease |
| rs115469976 | 6 | | | |
| rs3892710 | 6 | | Inflammation, Zellweger Syndrome, Coronary Artery Disease | |
| rs9275334 | 6 | | | |
| rs73405471 | 6 | HLA-DQA1 | | Esophageal Achalasia, Celiac Disease, Diabetes Mellitus, Insulin-Dependent |
| rs28371206 | 6 | | | |
| rs3135335 | 6 | | | |
| rs3135006 | 6 | | Brain Stem Infarctions, N syndrome | |
| rs3957146 | 6 | | Diabetes Mellitus | |
| rs3842752 | 11 | INS | Necrosis, Xeroderma Pigmentosum, Breast Neoplasms | Diabetes Mellitus, Permanent Neonatal, Maturity Onset Diabetes Mellitus In Young, Diabetes Mellitus |
| rs17555038 | 5 | SGCD | | Limb-Girdle Muscular Dystrophy Type 2F, Cardiomyopathy, Dilated, 1L, Cardiomyopathies |
| rs9273088 | 6 | HLA-DQA1 | | Esophageal Achalasia, Celiac Disease, Diabetes Mellitus, Insulin-Dependent |
| rs9273363 | 6 | HLA-DQB1-AS1 | Diabetes Mellitus, Arthritis, Rheumatoid, Autoimmune Diseases | Chronic Obstructive Airway Disease |
| rs9275373 | 6 | | Parkinson Disease, Isobutyryl-CoA dehydrogenase deficiency | |
| rs3135395 | 6 | | | |
| rs3104413 | 6 | | Autoimmune Diseases | |
| rs9275356 | 6 | | | |

**Supplementary Table 4.** Cross-reference of the 20 most highly activated SNPs, on average across multiple training runs, and known variant and gene associations for the GLN models trained on type 1 diabetes. The top SNPs are chosen according to the average of highest activated SNPs across 10 runs with different random seeds each. The top 3 LitVar values were selected according to the most co-occuring disease entities. For DisGeNET, the top three scoring gene-disease DisGeNET association scores were used.

| rsID | Chr. | Gene | Top 3 LitVar | Top 3 DisGeNET |
|---|---|---|---|---|
| rs3731239 | 9 | CDKN2A | Breast Neoplasms, Neoplasms, N syndrome | Esophageal Neoplasms, Lung Neoplasms, Melanoma |
| rs17017475 | 1 | | | |
| rs2427221 | 20 | CDH4 | | Longevity, Sarcoidosis, Polysomnography |
| rs12405608 | 1 | | | |
| rs34410881 | 1 | | | |
| rs2858584 | 22 | | | |
| rs7355020 | 1 | NAV1 | | Schizophrenia, Body Height, Cardiovascular Diseases |
| rs1043811 | 12 | RBM19 | | Diabetes Mellitus, Non-Insulin-Dependent, Response To Simvastatin, Blepharoptosis |
| rs10871782 | 18 | | | |
| rs75602167 | 22 | TOP3B | Alzheimer Disease | Schizophrenia, Autistic Disorder, Impaired Cognition |
| rs4977574 | 9 | CDKN2B-AS1 | Coronary Disease, Coronary Artery Disease, Myocardial Infarction | Endometriosis, Glaucoma, Open-Angle, Nasopharyngeal Carcinoma |
| rs2549505 | 16 | MAF | | Cataract 21, Multiple Types, Cataracts, Congenital, With Sensorineural Deafness, Down Syndrome-Like Facial Appearance, Short Stature, And Mental Retardation, Cataract, Pulverulent, Juvenile-Onset |
| rs17705635 | 18 | CHST9 | | Malignant Neoplasm Of Breast, Adolescent Idiopathic Scoliosis, Breast Carcinoma |
| rs6128184 | 20 | | | |
| rs8005039 | 14 | AKAP6 | | Atrial Fibrillation, Intellectual Disability, Malignant Neoplasm Of Breast |
| rs72699511 | 1 | | | |
| rs10784085 | 12 | | | |
| rs149144163 | 20 | | | |
| rs3184504 | 12 | SH2B3 | Diabetes Mellitus, Arthritis, Rheumatoid, Celiac Disease | Precursor Cell Lymphoblastic Leukemia Lymphoma, Thrombocythemia, Essential, Diabetes Mellitus, Insulin-Dependent |
| rs71331632 | 21 | | | |

**Supplementary Table 5.** Cross-reference of the 20 most highly activated SNPs and known variant and gene associations for the GLN model trained on acute myocardial infarction. The top 3 LitVar values were selected according to the most co-occuring disease entities. For DisGeNET, the top three scoring gene-disease DisGeNET association scores were used.

| rsID | Chr. | Gene | Top 3 LitVar | Top 3 DisGeNET |
|---|---|---|---|---|
| rs10975479 | 9 | | | |
| rs340921 | 9 | | | |
| rs8056488 | 16 | | | |
| rs71430382 | 2 | D2HGDH | | D-2-Hydroxyglutaric Aciduria 1, Combined D-2- And L-2-Hydroxyglutaric Aciduria, Amino Acid Metabolism, Inborn Errors |
| rs17293632 | 15 | SMAD3 | Crohn Disease, Inflammatory Bowel Diseases, Arthritis, Rheumatoid | Loeys-Dietz Syndrome 3, Colorectal Carcinoma, Loeys-Dietz Syndrome |
| rs2160203 | 2 | IL1RL1 | Diabetes Mellitus, Asthma, Rhinitis | Asthma, Arthritis, Adjuvant-Induced, Acute Lung Injury |
| rs343476 | 9 | | | |
| rs72777284 | 2 | | | |
| rs111543205 | 2 | D2HGDH | | D-2-Hydroxyglutaric Aciduria 1, Combined D-2- And L-2-Hydroxyglutaric Aciduria, Amino Acid Metabolism, Inborn Errors |
| rs2589559 | 10 | | | |
| rs167769 | 12 | STAT6 | Asthma, Alzheimer Disease, Pain | Solitary Fibrous Tumor, Asthma, Dermatitis, Atopic |
| Affx-37000939 | 5 | | | |
| rs9050 | 1 | TCHH | Alzheimer Disease, Properdin deficiency, X-linked | Uncombable Hair Syndrome, Dermatitis, Atopic, Curly Hair (Finding) |
| rs11642659 | 16 | LCMT1 | | Blastocyst Disintegration, Embryo Resorption, Embryo Death |
| rs2706347 | 5 | RAD50 | Asthma, Dermatitis, Atopic, Rhinitis | Nijmegen Breakage Syndrome-Like Disorder, Asthma, Breast Cancer, Familial |
| rs870301 | 2 | BOK-AS1 | | Forced Expiratory Volume Function, Inflammatory Bowel Diseases, Vital Capacity |
| rs17612633 | 6 | HLA-DQA1 | | Esophageal Achalasia, Celiac Disease, Diabetes Mellitus, Insulin-Dependent |
| Affx-8225028 | 12 | | | |
| rs62298921 | 3 | | | |
| rs3771180 | 2 | IL1RL1 | Asthma, Rhinitis, Allergic, Seasonal, Obesity | Asthma, Arthritis, Adjuvant-Induced, Acute Lung Injury |

**Supplementary Table 6.** Cross-reference of the 20 most highly activated SNPs and known variant and gene associations for the GLN model trained on asthma. The top 3 LitVar values were selected according to the most co-occuring disease entities. For DisGeNET, the top three scoring gene-disease DisGeNET association scores were used.

| rsID | Chr. | Gene | Top 3 LitVar | Top 3 DisGeNET |
|---|---|---|---|---|
| rs17042171 | 4 | | Atrial Fibrillation, Diabetes Mellitus, Hypertension | |
| rs3853445 | 4 | | Atrial Fibrillation, Hypertension, Properdin deficiency, X-linked | |
| rs17042081 | 4 | | | |
| rs13141190 | 4 | | | |
| rs10033464 | 4 | | Atrial Fibrillation, Stroke, Cerebral Infarction | |
| rs1448817 | 4 | | Atrial Fibrillation | |
| rs4074536 | 1 | CASQ2 | Tachycardia, Ventricular, Long QT Syndrome, Atrial Fibrillation | Stress-Induced Polymorphic Ventricular Tachycardia, Ventricular Tachycardia, Catecholaminergic Polymorphic, 1 (Disorder), Tachycardia, Ventricular |
| rs1906610 | 4 | | | |
| rs13376333 | 1 | KCNN3 | Atrial Fibrillation, Hypertension, Diabetes Mellitus | Zimmerman Laband Syndrome, Atrial Fibrillation, Schizophrenia |
| rs7667461 | 4 | | | |
| rs521511 | 4 | | | |
| rs72811957 | 17 | NTN1 | | Mirror Movements 4, Subarachnoid Hemorrhage, Cerebral Edema |
| rs117984853 | 6 | | Atrial Fibrillation | |
| rs10908444 | 1 | KCNN3 | | Zimmerman Laband Syndrome, Atrial Fibrillation, Schizophrenia |
| rs1386389 | 4 | | | |
| rs17825726 | 22 | | | |
| rs5765546 | 22 | | | |
| rs4845695 | 1 | PMVK | Drug-Related Side Effects and Adverse Reactions, 13q deletion syndrome | Porokeratosis Of Mibelli, Porokeratosis, Linear, Porokeratosis |
| rs2106261 | 16 | ZFHX3 | Atrial Fibrillation, Hypertension, Inflammation | Atrial Fibrillation, Cerebrovascular Accident, Malignant Neoplasm Of Prostate |
| rs5886821 | 7 | LOC107986838 | | |

**Supplementary Table 7.** Cross-reference of the 20 most highly activated SNPs and known variant and gene associations for the GLN model trained on atrial fibrillation and flutter. The top 3 LitVar values were selected according to the most co-occuring disease entities. For DisGeNET, the top three scoring gene-disease DisGeNET association scores were used.

| rsID | Chr. | Gene | Top 3 LitVar | Top 3 DisGeNET |
|---|---|---|---|---|
| rs117888135 | 10 | | | |
| rs12940887 | 17 | ZNF652 | Arthritis, Rheumatoid, Sveinsson Chorioretinal Atrophy, Ataxia Telangiectasia | Malignant Neoplasms, Malignant Neoplasm Of Prostate, Prostate Carcinoma |
| rs34328549 | 19 | INSR | | Donohue Syndrome, Rabson-Mendenhall Syndrome, Diabetes Mellitus, Non-Insulin-Dependent |
| rs4932370 | 15 | | Schmid-Fraccaro syndrome, Ataxia Telangiectasia | |
| rs9647448 | 4 | LCORL | | Birth Weight, Body Height, Cardiovascular Diseases |
| rs4987082 | 17 | PHB | Autistic Disorder | Malignant Neoplasm Of Breast, Mammary Neoplasms, Experimental, Breast Carcinoma |
| rs160889 | 5 | | | |
| rs16834635 | 1 | PBX1 | | Burkitt Lymphoma, Congenital Anomalies Of Kidney And Urinary Tract Syndrome With Or Without Hearing Loss, Abnormal Ears, Or Developmental Delay, Precursor B-Cell Lymphoblastic Leukemia |
| rs4766897 | 12 | ACAD10 | | Age Related Macular Degeneration, Alcohol Consumption, Coronary Heart Disease |
| rs6848130 | 4 | | | |
| rs52824916 | 12 | FAM186B | | Nephronophthisis, Systolic Pressure |
| rs2963446 | 5 | | | |
| rs356986 | 2 | | | |
| rs35208507 | 16 | PDILT | | Colorectal Carcinoma, Blood Urea Nitrogen Measurement, Cardiovascular Diseases |
| rs35112858 | 8 | MSRA | | Schizophrenia, Drug Abuse, Drug Habituation |
| rs7354757 | 21 | MORC3 | | Body Height, White Blood Cell Count Procedure, Diastolic Blood Pressure |
| rs10004996 | 4 | LINC02513 | | Lymphocyte Count Measurement |
| rs7561317 | 2 | | Obesity, Diabetes Mellitus, Type 2, Diabetes Mellitus | |
| rs13115333 | 4 | | | |
| rs1054707 | 4 | BDH2 | | Neoplasms, Iron Deficiency, Tumor Cell Invasion |

**Supplementary Table 8.** Cross-reference of the 20 most highly activated SNPs and known variant and gene associations for the GLN model trained on hypertension. The top 3 LitVar values were selected according to the most co-occuring disease entities. For DisGeNET, the top three scoring gene-disease DisGeNET association scores were used.

| rsID | Chr. | Gene | Top 3 LitVar | Top 3 DisGeNET |
|---|---|---|---|---|
| rs733175 | 4 | | Alzheimer Disease, Parkinson Disease, Nephrolithiasis | |
| rs2231142 | 4 | ABCG2 | Gout, Hyperuricemia, Neoplasms | Uric Acid Concentration, Serum, Quantitative Trait Locus 1, Hyperuricemia, Gout |
| rs4148157 | 4 | ABCG2 | Brain Neoplasms, Properdin deficiency, X-linked | Uric Acid Concentration, Serum, Quantitative Trait Locus 1, Hyperuricemia, Gout |
| rs7671266 | 4 | | Ataxia Telangiectasia | |
| rs6855911 | 4 | SLC2A9 | Anxiety Disorders, Phobic Disorders, Metabolic Diseases | Hypouricemia, Renal, 2, Renal Hypouricemia, Hyperuricemia |
| rs13129697 | 4 | SLC2A9 | Diabetes Mellitus, Kidney Diseases, Hypertension | Hypouricemia, Renal, 2, Renal Hypouricemia, Hyperuricemia |
| rs737267 | 4 | SLC2A9 | Glycogen Storage Disease Type V, Parkinson Disease, Hypertension | Hypouricemia, Renal, 2, Renal Hypouricemia, Hyperuricemia |
| rs4698023 | 4 | | | |
| rs4481233 | 4 | SLC2A9 | Familial benign hypercalcemia, type 3, Gout | Hypouricemia, Renal, 2, Renal Hypouricemia, Hyperuricemia |
| rs73203233 | 21 | | | |
| rs16890979 | 4 | SLC2A9 | Gout, Zellweger Syndrome, Nephrolithiasis | Hypouricemia, Renal, 2, Renal Hypouricemia, Hyperuricemia |
| rs7671500 | 4 | | | |
| rs11048289 | 12 | | | |
| rs6834555 | 4 | | Alzheimer Disease, Psychoses, Substance-Induced, Properdin deficiency, X-linked | |
| rs7442295 | 4 | SLC2A9 | Glycogen Storage Disease Type V, Myocardial Ischemia, Gout | Hypouricemia, Renal, 2, Renal Hypouricemia, Hyperuricemia |
| rs995014 | 21 | N6AMT1 | | Glomerular Filtration Rate, Malignant Neoplasms, Primary Malignant Neoplasm |
| rs4809565 | 20 | KCNQ2 | | Seizures, Benign Familial Neonatal, 1, Epileptic Encephalopathy, Early Infantile, 7, Familial Benign Neonatal Epilepsy |
| rs4910001 | 11 | GALNT18 | | Rheumatoid Arthritis, Serum Albumin Measurement, Gastric Adenocarcinoma |
| rs3114020 | 4 | ABCG2 | Carcinoma, Non-Small-Cell Lung, Properdin deficiency, X-linked, Adenocarcinoma | Uric Acid Concentration, Serum, Quantitative Trait Locus 1, Hyperuricemia, Gout |
| rs3760627 | 19 | CLPTM1 | Alzheimer Disease | Pancreatic Neoplasm, Malignant Neoplasm Of Pancreas, Alzheimer'S Disease |

**Supplementary Table 9.** Cross-reference of the 20 most highly activated SNPs and known variant and gene associations for the GLN model trained on gout. The top 3 LitVar values were selected according to the most co-occuring disease entities. For DisGeNET, the top three scoring gene-disease DisGeNET association scores were used.

| rsID | Chr. | Gene | Top 3 LitVar | Top 3 DisGeNET |
|---|---|---|---|---|
| rs4506565 | 10 | TCF7L2 | Diabetes Mellitus, Diabetes Mellitus, Type 2, Obesity | Colorectal Carcinoma, Diabetes Mellitus, Non-Insulin-Dependent, Colorectal Neoplasms |
| rs12243326 | 10 | TCF7L2 | Polycystic Ovary Syndrome, Diabetes Mellitus, Hypoglycemia | Colorectal Carcinoma, Diabetes Mellitus, Non-Insulin-Dependent, Colorectal Neoplasms |
| rs12255372 | 10 | TCF7L2 | Diabetes Mellitus, Diabetes Mellitus, Type 2, Obesity | Colorectal Carcinoma, Diabetes Mellitus, Non-Insulin-Dependent, Colorectal Neoplasms |
| rs11196175 | 10 | TCF7L2 | Metabolic Diseases, Choroideremia, Ovarian Neoplasms | Colorectal Carcinoma, Diabetes Mellitus, Non-Insulin-Dependent, Colorectal Neoplasms |
| rs7903146 | 10 | TCF7L2 | Diabetes Mellitus, Diabetes Mellitus, Type 2, Obesity | Colorectal Carcinoma, Diabetes Mellitus, Non-Insulin-Dependent, Colorectal Neoplasms |
| rs2796441 | 9 | LOC101927502 | Diabetes Mellitus, Diabetes Mellitus, Type 2, Ataxia Telangiectasia | |
| rs62208714 | 20 | PTPRT | | Leukemia, Myelocytic, Acute, Colorectal Carcinoma, Malignant Neoplasm Of Lung |
| rs11257655 | 10 | | Properdin deficiency, X-linked, Diabetes Mellitus, Diabetes Mellitus, Type 2 | |
| rs7069060 | 10 | | Retinoschisis | |
| rs7018475 | 9 | | Diabetes Mellitus, Obesity, Histidinemia | |
| rs9273363 | 6 | HLA-DQB1-AS1 | Diabetes Mellitus, Arthritis, Rheumatoid, Autoimmune Diseases | Chronic Obstructive Airway Disease |
| rs163177 | 11 | KCNQ1 | Properdin deficiency, X-linked, Coronary Artery Disease | Jervell-Lange Nielsen Syndrome, Long Qt Syndrome 1, Short Qt Syndrome 2 (Disorder) |
| rs17817449 | 16 | FTO | Obesity, Breast Neoplasms, Diabetes Mellitus | Growth Retardation, Developmental Delay, Coarse Facies, And Early Death, Malignant Neoplasm Of Breast, Diabetes Mellitus, Non-Insulin-Dependent |
| rs388508 | 10 | LYZL2 | | Dental Caries |
| rs35368011 | 5 | C5orf67 | | Alcohol Consumption, Cardiovascular Diseases, Diabetes Mellitus, Non-Insulin-Dependent |
| rs12221133 | 10 | CDC123 | | Diabetes Mellitus, Non-Insulin-Dependent, Fibrosarcoma, Forced Expiratory Volume Function |
| rs2484892 | 9 | | | |
| rs62020600 | 16 | TMEM114 | | Cholelithiasis, Vital Capacity, Cholecystolithiasis |
| rs1046320 | 4 | WFS1 | Diabetes Mellitus, Type 2, Diabetes Mellitus | Wolfram Syndrome 1, Wolfram Syndrome, Wolfram-Like Syndrome, Autosomal Dominant |
| rs1572053 | 20 | EYA2 | | Alcohol Consumption, Diabetes Mellitus, Non-Insulin-Dependent, Eczema |

**Supplementary Table 10.** Cross-reference of the 20 most highly activated SNPs and known variant and gene associations for the GLN model trained on type 2 diabetes. The top 3 LitVar values were selected according to the most co-occuring disease entities. For DisGeNET, the top three scoring gene-disease DisGeNET association scores were used.

| rsID | Chr. | Gene | Top 3 LitVar | Top 3 DisGeNET |
|---|---|---|---|---|
| rs231727 | 2 | | Polyendocrinopathies, Autoimmune, 211750, Ataxia Telangiectasia | |
| rs3184504 | 12 | SH2B3 | Diabetes Mellitus, Arthritis, Rheumatoid, Celiac Disease | Precursor Cell Lymphoblastic Leukemia Lymphoma, Thrombocythemia, Essential, Diabetes Mellitus, Insulin-Dependent |
| rs17364832 | 13 | SPATA13 | | Autoimmune Diseases, Hypothyroidism, Respiratory Function Tests |
| rs653178 | 12 | ATXN2 | Arthritis, Rheumatoid, Diabetes Mellitus, Celiac Disease | Spinocerebellar Ataxia Type 2, Amyotrophic Lateral Sclerosis, Parkinson Disease, Late-Onset |
| rs10028213 | 4 | LOC107986195 | Hypothyroidism, Thyroid Neoplasms | |
| rs1348386 | 9 | PTCSC2 | | Thyroid Carcinoma, Alopecia Areata, Autoimmune Diseases |
| rs7323885 | 13 | SPATA13 | | Autoimmune Diseases, Hypothyroidism, Respiratory Function Tests |
| rs2412970 | 22 | HORMAD2 | Colitis, Crohn Disease, Isobutyryl-CoA dehydrogenase deficiency | Malignant Neoplasm Of Lung, Iga Glomerulonephritis, Ulcerative Colitis |
| rs925489 | 9 | PTCSC2 | Thyroid cancer, papillary, Autoimmune Diseases, Hypertension | Thyroid Carcinoma, Alopecia Areata, Autoimmune Diseases |
| rs310405 | 6 | | Meige Syndrome, Charcot-Marie-Tooth disease, Type 1D | |
| rs2412971 | 22 | HORMAD2 | Inflammatory Bowel Diseases, Isobutyryl-CoA dehydrogenase deficiency, Proteinuria | Malignant Neoplasm Of Lung, Iga Glomerulonephritis, Ulcerative Colitis |
| rs6679677 | 1 | PHTF1 | Arthritis, Rheumatoid, Diabetes Mellitus, Lupus Erythematosus, Systemic | Rheumatoid Arthritis, Diabetes Mellitus, Insulin-Dependent |
| rs7574865 | 2 | STAT4 | Lupus Erythematosus, Systemic, Arthritis, Rheumatoid, Autoimmune Diseases | Lupus Erythematosus, Systemic, Behcet Syndrome, Rheumatoid Arthritis |
| rs12575636 | 11 | | Gaucher Disease | |
| rs654537 | 6 | BACH2 | | Crohn Disease, Diabetes Mellitus, Insulin-Dependent, Celiac Disease |
| rs1024161 | 2 | | Graves Disease, Diabetes Mellitus, Diabetes Mellitus, Type 1 | |
| rs10452226 | 4 | LOC105377483 | | |
| rs118173218 | 14 | ESRRB | | Deafness, Autosomal Recessive 35, Hearing Impairment, Nonsyndromic Deafness |
| rs2233955 | 6 | C6orf15 | | Sarcoidosis, Human Immunodeficiency Virus Type 1, Susceptibility To, Hiv-1, Resistance To |
| rs4818324 | 21 | | | |

**Supplementary Table 11.** Cross-reference of the 20 most highly activated SNPs and known variant and gene associations for the GLN model trained on hypothyroidism. The top 3 LitVar values were selected according to the most co-occuring disease entities. For DisGeNET, the top three scoring gene-disease DisGeNET association scores were used.

| rsID | Chr. | Gene | Top 3 LitVar | Top 3 DisGeNET |
|---|---|---|---|---|
| rs12255372 | 10 | TCF7L2 | Diabetes Mellitus, Diabetes Mellitus, Type 2, Obesity | Colorectal Carcinoma, Diabetes Mellitus, Non-Insulin-Dependent, Colorectal Neoplasms |
| rs7903146 | 10 | TCF7L2 | Diabetes Mellitus, Diabetes Mellitus, Type 2, Obesity | Colorectal Carcinoma, Diabetes Mellitus, Non-Insulin-Dependent, Colorectal Neoplasms |
| rs76769781 | 20 | | | |
| rs117697300 | 21 | | | |
| rs12243326 | 10 | TCF7L2 | Polycystic Ovary Syndrome, Diabetes Mellitus, Hypoglycemia | Colorectal Carcinoma, Diabetes Mellitus, Non-Insulin-Dependent, Colorectal Neoplasms |
| rs1880008 | 22 | | | |
| rs4506565 | 10 | TCF7L2 | Diabetes Mellitus, Diabetes Mellitus, Type 2, Obesity | Colorectal Carcinoma, Diabetes Mellitus, Non-Insulin-Dependent, Colorectal Neoplasms |
| rs4751323 | 10 | DOCK1 | | Drug Abuse, Drug Habituation, Drug Use Disorders |
| rs7752155 | 6 | | | |
| rs6882745 | 5 | | | |
| rs251404 | 5 | PIK3R1 | | Short Syndrome, Insulin Resistance, Agammaglobulinemia 7, Autosomal Recessive |
| rs13124487 | 4 | PPP2R2C | | Bipolar Disorder, Pathological Accumulation Of Air In Tissues, Diabetes Mellitus, Non-Insulin-Dependent |
| rs17065659 | 5 | | | |
| rs4431094 | 3 | | | |
| rs713928 | 22 | TAFA5 | | Pancreatic Carcinoma, Malignant Neoplasm Of Pancreas, Adenovirus Infections |
| rs2869310 | 20 | | | |
| rs1106440 | 7 | CNTNAP2 | | Pitt-Hopkins-Like Syndrome 1, Cortical Dysplasia With Focal Epilepsy Syndrome, Autism Spectrum Disorders |
| rs3016382 | 11 | OPCML | | Ovarian Neoplasm, Malignant Neoplasm Of Ovary, Schizophrenia |
| rs11176379 | 12 | GRIP1 | | Cryptophthalmos Syndrome, Fraser Syndrome 3, Schizophrenia |
| rs79768529 | 4 | LOC105377567 | | |

**Supplementary Table 12.** Cross-reference of the 20 most highly activated SNPs and known variant and gene associations for the GLN model trained on type 2 diabetes, when including clinical and biochemical measurements. The top 3 LitVar values were selected according to the most co-occuring disease entities. For DisGeNET, the top three scoring gene-disease DisGeNET association scores were used.

| rsID | Chr. | Gene | Top 3 LitVar | Top 3 DisGeNET |
|---|---|---|---|---|
| rs532965 | 6 | | | |
| rs6931277 | 6 | | Alzheimer Disease | |
| rs3104413 | 6 | | Autoimmune Diseases | |
| rs3830127 | 6 | HLA-DRB1 | | Crohn Disease, Lupus Erythematosus, Systemic, Narcolepsy |
| rs34250758 | 6 | HLA-DQA1 | | Esophageal Achalasia, Celiac Disease, Diabetes Mellitus, Insulin-Dependent |
| rs2760976 | 6 | | | |
| rs34855541 | 6 | | | |
| rs521539 | 6 | | | |
| rs2858333 | 6 | | Testicular Neoplasms | |
| rs660895 | 6 | | Parkinson Disease, Arthritis, Rheumatoid, Lupus Erythematosus, Systemic | |
| rs2647087 | 6 | | Pancreatitis, Colonic Diseases, Testicular Neoplasms | |
| rs7745656 | 6 | | Pancreatitis | |
| rs35265698 | 6 | | | |
| rs9268515 | 6 | | Parkinson Disease, Properdin deficiency, X-linked | |
| rs9275555 | 6 | | | |
| rs5004277 | 6 | | | |
| rs3793127 | 6 | BTNL2 | 211750, Ataxia Telangiectasia | Sarcoidosis, Berylliosis, Beryllium Disease |
| Affx-28502467 | 6 | | | |
| rs3104415 | 6 | | | |
| rs111586361 | 6 | HLA-DRB6 | | Child Development Disorders, Pervasive, Lymphoma, Follicular, Multiple Sclerosis |

**Supplementary Table 13.** Cross-reference of the 20 most highly activated SNPs and known variant and gene associations for the LASSO model trained on rheumatoid arthritis. The top 3 LitVar values were selected according to the most co-occuring disease entities. For DisGeNET, the top three scoring gene-disease DisGeNET association scores were used.

| rsID | Chr. | Gene | Top 3 LitVar | Top 3 DisGeNET |
|---|---|---|---|---|
| rs474235 | 4 | NSD2 | | Wolf-Hirschhorn Syndrome, Pitt-Rogers-Danks Syndrome, Microcephaly |
| rs55994383 | 15 | | | |
| rs2395163 | 6 | | Parkinson Disease, Schizophrenia, Parkinson Disease, Secondary | |
| rs4309342 | 15 | | | |
| rs7600206 | 2 | PAX3 | | Alveolar Rhabdomyosarcoma, Waardenburg Syndrome Type 1, Waardenburg Syndrome |
| rs115079157 | 2 | | | |
| rs3830127 | 6 | HLA-DRB1 | | Crohn Disease, Lupus Erythematosus, Systemic, Narcolepsy |
| rs117290280 | 16 | RAB11FIP3 | | Platelet Count Measurement, Mean Corpuscular Volume (Result), Finding Of Mean Corpuscular Hemoglobin |
| rs1273150 | 14 | DAAM1 | | Body Height, Tumor Cell Invasion, Malignant Neoplasm Of Breast |
| rs75021444 | 8 | | | |
| rs9928176 | 16 | | | |
| rs660895 | 6 | | Parkinson Disease, Arthritis, Rheumatoid, Lupus Erythematosus, Systemic | |
| rs9272226 | 6 | | Lupus Erythematosus, Systemic | |
| rs262929 | 6 | | | |
| rs2179685 | 20 | | | |
| rs62033174 | 16 | | | |
| rs9268645 | 6 | HLA-DRA | Diabetes Mellitus, Lupus Erythematosus, Systemic, Arthritis, Rheumatoid | Parkinson Disease, Multiple Sclerosis, Alcoholic Intoxication, Chronic |
| rs73248761 | 13 | | | |
| rs1320591 | 1 | | | |
| rs12044963 | 1 | KCND3 | Atrial Fibrillation, Brain Stem Infarctions | Spinocerebellar Ataxia 19, Brugada Syndrome 9, Brugada Syndrome (Disorder) |

**Supplementary Table 14.** Cross-reference of the 20 most highly activated SNPs and known variant and gene associations for the GLN model trained on rheumatoid arthritis. The top 3 LitVar values were selected according to the most co-occuring disease entities. For DisGeNET, the top three scoring gene-disease DisGeNET association scores were used.

| Allele | $\beta$ | Std. Err. | Z | $P > \|z\|$ | 0.025 CI | 0.975 CI | Odds |
|---|---|---|---|---|---|---|---|
| rs9273363 CC | -2.671 | 0.125 | -21.43 | $6.38 \times 10^{-102}$ | -2.915 | -2.426 | 0.06921 |
| rs9273363 CA | -2.267 | 0.124 | -18.31 | $6.771 \times 10^{-75}$ | -2.509 | -2.024 | 0.1036 |
| rs9273363 AA | -1.198 | 0.126 | -9.469 | $2.818 \times 10^{-21}$ | -1.445 | -0.95 | 0.3019 |
| rs3842752 GG | -2.487 | 0.123 | -20.3 | $1.334 \times 10^{-91}$ | -2.727 | -2.247 | 0.08316 |
| rs3842752 GA | -2.863 | 0.126 | -22.81 | $3.936 \times 10^{-115}$ | -3.109 | -2.617 | 0.0571 |
| rs3842752 AA | -2.849 | 0.151 | -18.82 | $5.622 \times 10^{-79}$ | -3.145 | -2.552 | 0.05792 |

**Supplementary Table 15.** Results of training a logistic regression model with SNPs rs9273363 and rs3842752 as inputs, encoded in a one-hot format, with type I diabetes diagnosis as the target variable. The odds ratios for rs9273363 (C→A) and rs3842752 (A→G) were 4.3620 and 1.407, respectively.
